# Tracking polygenic adaptation: phenotypic stasis after rapid adaptation in large populations of *Drosophila*

**DOI:** 10.1101/2025.06.09.658643

**Authors:** Claudia Ramirez-Lanzas, Marlies Dolezal, Sarah Signor, Christian Schlötterer, Neda Barghi

**Affiliations:** Institut Für Populationsgenetik, Vetmeduni Vienna, Vienna 1210, Austria; Vienna Graduate School of Population Genetics, Vetmeduni Vienna, Vienna 1210, Austria; Plattform Bioinformatik und Biostatistik, Vetmeduni Vienna, Vienna 1210, Austria; Department of Biological Sciences, North Dakota State University, Fargo, North Dakota 581042; Max Planck Institute for Evolutionary Biology, Plön 24306, Germany

## Abstract

Most adaptive traits have a polygenic basis, and their evolution in response to environmental change results from the cumulative effect of many underlying loci. Theoretical studies predict two phases of phenotypic evolution: an initial directional phase, characterized by shifts in the mean of the trait towards a new optimum; and a subsequent stabilizing phase, in which the mean phenotype remains constant. However, there is limited empirical evidence to confirm these predictions. We exposed six replicate populations of *Drosophila simulans*, each with 100,000 individuals, to a new high-protein diet for 31 generations. Using time-series of high-order and molecular phenotypes, we characterized the phases of polygenic adaptation and inferred the transition between the directional and stabilizing selection phases empirically. The initial (directional) phase was characterized by a rapid phenotypic response, followed by a plateau after seven generations of adaptation, suggesting proximity to the new trait optimum (i.e. the stabilizing phase). The phenotypic responses were highly parallel among the replicate populations throughout the experiment. The contribution of pleiotropic loci to adaptive phenotypic evolution varied across the different phases of adaptation. Adaptation throughout the experiment was mainly facilitated by genes with low-to-intermediate pleiotropy, while highly pleiotropic genes played a significant role in the initial phase of adaptation. Parallel evolutionary responses across the replicate populations were also driven by genes with low-to-intermediate pleiotropy. Our findings support the predictions of theoretical models and provide empirical evidence for directional and stabilizing phases of polygenic adaptation.

## Introduction

Most traits under natural selection are quantitative, with a continuous and typically normal distribution (1). The collective findings of many QTL mapping, GWAS, and evolution studies in species ranging from *Drosophila* (2) to humans (3) have shown that these quantitative traits are complex and determined by many underlying loci. Most quantitative traits are under stabilizing selection (1, 3) which stabilizes the population’s mean on an optimal trait value, known as trait optimum. After a sudden environmental change which shifts the trait’s optimum, adaptation would occur through the collective, often small, frequency changes across many alleles, i.e. polygenic adaptation. Theoretical (4, 5) and simulation studies (6–9) predict that adaptation of quantitative traits has at least two phases, each with distinct genomic and phenotypic characteristics. Right after a shift in optimum, due to the frequency change of adaptive alleles the population’s mean phenotype changes toward the new optimum. This initial phase of adaptation is rapid, resembling directional selection (4, 6, 7). In the second phase, after the population’s mean phenotype approaches the new optimum, the combined effects of stabilizing selection and genetic drift result in the fixation and loss of alleles (alleles with effects aligned with the shift, as well as alleles with opposing effects, could be fixed or lost, see (4)) while the phenotype’s mean phenotype stabilizes on the new optimum.

Empirical data supporting theoretical predictions remain limited for several reasons. First, most theoretical models assume a stable environment after an initial shift in trait optimum, allowing stabilizing selection to act. However, natural populations often experience seasonal fluctuations which contradicts this assumption. Second, the trait under selection and its new optimum are often unknown in most (experimental and natural) populations. These factors hinder our ability to determine when the populations reach the new optimum and to distinguish between directional and stabilizing phases of adaptation. The predicted phases of adaptation (4–9) suggest that methods for the detection of adaptation should incorporate the temporal phenotypic and genomic dynamics. Additionally, to align with the model assumptions, a stable environment should follow the initial change. Experimental evolution provides an ideal framework for testing the theoretical predictions by allowing the collection of temporal data in controlled and constant environments across replicate populations (6).

Predictions of allele frequencies changes during complex traits adaptation depend on trait polygenicity, allele frequency and effect size, and the magnitude of trait optimum shift. Theoretical studies have assumed different optimum shifts (4, 8) and trait polygenicity (4, 9). Despite differing genomic predictions, a consistent pattern across all studies is that the phenotypic mean plateaus as populations approach the new optimum. Simulations (6, 7) show that temporal phenotypic patterns can distinguish a transition from directional to stabilizing phases of adaptation. Identifying this transition helps pinpoint when populations approach the new optimum, even without knowing the selected trait.

To characterize the temporal patterns of phenotypic changes after a shift in the trait optimum we exposed unprecedentedly large populations of *Drosophila simulans* to a novel selection regime with a high-protein diet in the laboratory for 40 non-overlapping generations. We quantified phenotypic changes at multiple time points in evolving populations by collecting data across different levels of the biological hierarchy, including molecular traits (gene expression) and high-order phenotypes (lipid content, weight, and fecundity, as a fitness component). We characterized the temporal patterns of adaptive phenotypic change and identified the transition from the directional phase to the stabilizing selection phase. This is the first empirical test of polygenic adaptation models showing rapid adaptation followed by phenotypic stasis.

## Results

We performed round-robin crosses using 93 *Drosophila simulans* inbred lines, maintained them for 15 generations, and established a founder population by combining equal proportions of these crosses (Fig. S1). Six replicate populations each with 100,000 individuals were set up from the founder population and exposed to a novel selection regime that consists of a high-protein (protein:carbohydrate ratio 1:1) diet (Fig. S2, Table S1) for 40 non-overlapping generations (Fig. S1). Populations were exposed to a complex selection regime consisting of multiple selection pressures, including a new high-protein diet and maintenance at 25 °C with a 12-days developmental time. Adaptation to high-protein diet (10), temperature (11, 12) and developmental time (13) are all polygenic traits. Thus, this complex selection regime changed the optimal phenotype of the populations, i.e. trait optimum, for multiple polygenic traits.

## Rapid change of high-order and molecular phenotypes

To determine the phenotypic changes as populations evolve on the high-protein diet, we measured the phenotypic status of the founder population and that of the evolving populations after 7 and 31 generations. Each population was maintained in a common garden for 2 generations on high-protein diet with controlled density before phenotyping to reduce parental effects (Fig. S1). This assured that the phenotypic differences observed between the founder and evolved populations reflected adaptation to the new environment, not transgenerational plasticity. First, we assessed the adaptive response of the evolved populations by measuring female fecundity, which is a proxy of fitness. The evolved populations at generations 7 and 31 were significantly more fecund, hence fitter, than the founder population (Fig. 1A). Weight, a high-order phenotype, also showed a rapid response, with significant decreases in weight observed in both females and males after only 7 generations (Fig. S3). The weight of both females and males increased significantly after generation 7.

**Fig. 1.**
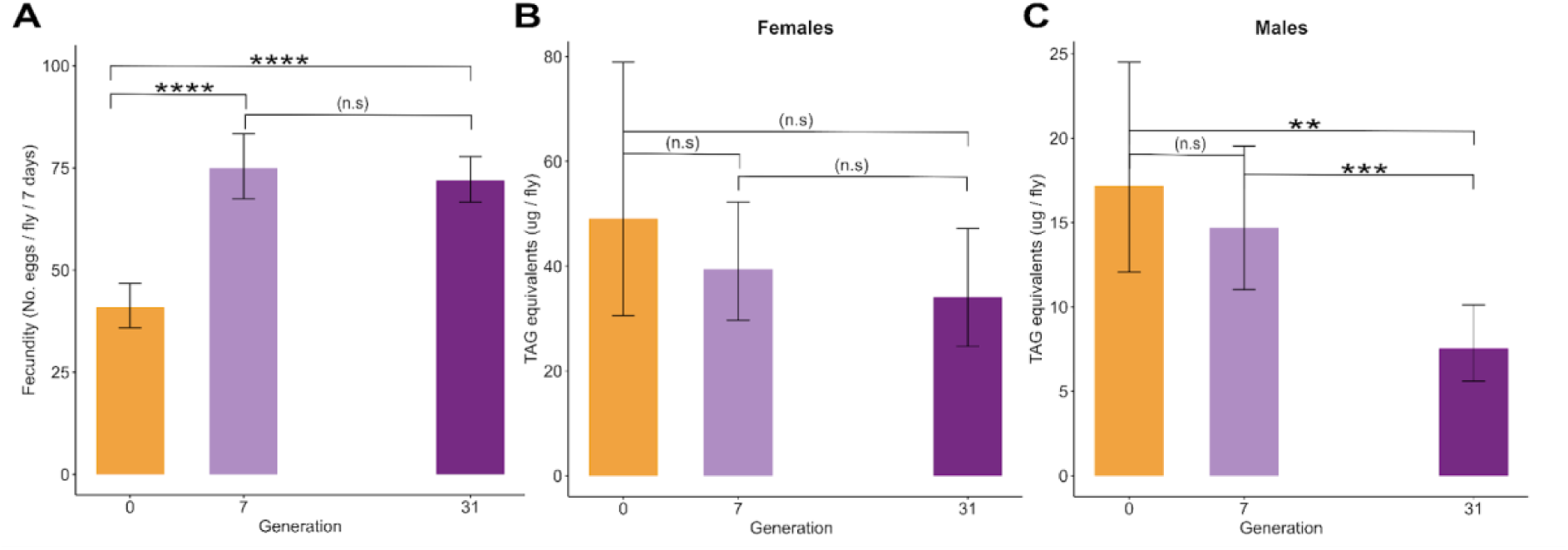
Adaptive response of high-order phenotypes, fecundity (A) and lipid content (B,C). A) Females from the evolved populations showed significantly higher fecundity at generations 7 and 31 compared to the founder population. Fecundity plateaued after generation 7, with no significant difference between generations 7 and 31. Fecundity is presented as the average number of eggs laid by a female during 7 days. B,C) In males, the amount of TAG was significantly less than in the founder population after 31 generations but not at generation 7. The lipid content in females from the evolved populations was not significantly different from the founder population. Asterisks indicate significance levels: *p*-value < 0.01 (**), *p*-value < 0.001 (***), and *p*-value < 0.0001 (****). Bars depict the predicted means of the linear mixed model, and error bars show the 95% confidence intervals of the least-square means.

As many of the genetic variants underlying complex traits are located in regulatory regions (14, 15), adaptation may manifest through changes in the expression of adaptive genes. Therefore, we investigated changes in the gene expression as the underlying molecular basis of high-order phenotypic changes. We measured the gene expression of males in the founder population and in the evolved populations at generations 7 and 31, after maintenance in a common garden on high-protein diet for 2 generations. Principal component analysis (PCA) of global transcriptomic variation among 10,197 expressed genes revealed clear differentiation between the founder population and evolved populations (PC1, 26.19%, Fig. S4A). However, the expression variation which distinguishes the evolved populations at generation 7 and 31 is small (PC3, 7.89%, Fig S4B). A total of 6,354 genes had significant changes in their expression in the evolved populations when compared to the founder population. Of these, a total of 4,867 genes (76.6%) were shared between generations 7 and 31 (Fig. 2). A fraction of the differentially expressed genes (710 genes, 11.17%) was identified only at generation 7 (‘complete-reverse’ group) while 777 genes (12.2%) showed a significant expression change only at generation 31 (‘late-response’ group, Fig. 2).

**Fig. 2.**
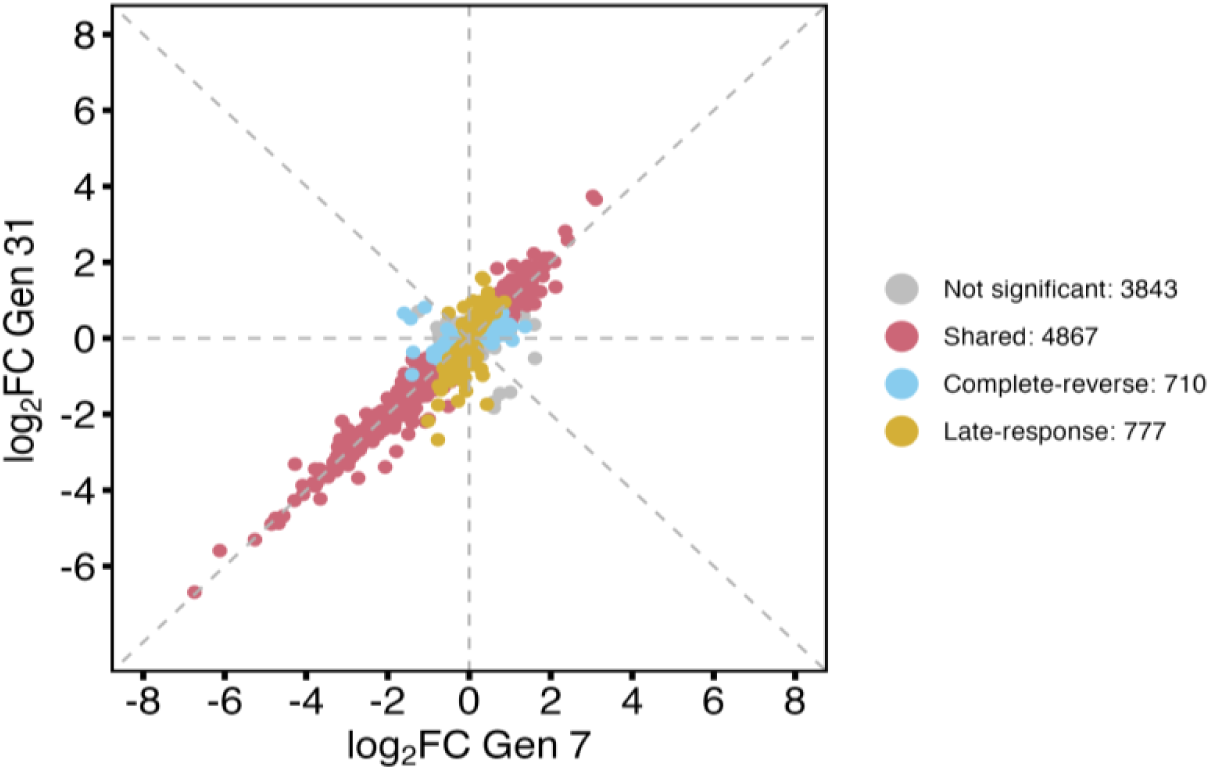
Highly correlated transcriptomic responses after 7 and 31 generations of adaptation. There was a high correlation in the transcriptomic response of the evolved populations at generation 7 and 31 (Pearson correlation coefficient of log_2_FC = 0.96). Transcriptomic response is expressed as the log_2_ fold-change (log_2_FC) between the founder and evolved populations at generation 7 (x axis) and generation 31 (y axis). ‘Shared’ group: identified differentially expressed genes (DEGs) at both generations 7 and 31, ‘complete-reverse’ group: DEGs that are identified only at generation 7, and ‘late-response’ group: DEGs that are identified only at generation 31.

To examine the functional associations of the differentially expressed genes (DEGs), we performed gene ontology (GO), pathway and tissue enrichment analyses (Table S2). DEGs were enriched in several GO terms, such as ‘proteolysis’ (GO:0006508), ‘siRNA processing’ (GO:0030422), ‘sexual reproduction’ (GO:0019953), and (GO:0015031) ‘protein transport’ (Table S2A, Fig. S5). Several pathways related to protein metabolism, e.g. ‘beta-alanine metabolism’ and ‘valine, leucine and isoleucine degradation’, ‘carbohydrate metabolism’, and ‘lipid metabolism’ were also enriched (Table S2B, Fig. S6). DEGs were also enriched in several tissues including fat body, midgut, malpighian tubule, heart and testis (Table S2C, Fig. S7). Collectively, the enrichment analyses showed a diverse response during adaptation to the selection environment involving many biological processes, tissues and pathways. This is expected given that the new environment required adaptation to a new diet, temperature regime and developmental time. In particular, enrichment of pathways related to protein metabolism and transport were expected as exposure to a high-protein diet was one of the main factors in the new selection regime.

## Stasis of high-order and molecular phenotypes

Theoretical models of polygenic adaptation (4) predict that after an environmental change, the population’s mean phenotype changes rapidly before slowing down to a pause when it nears the new optimum. We expect to observe stasis in phenotypes if the evolved populations are close to the optimal phenotype. At the high-order phenotype level, no significant difference was observed in the fecundity of evolved populations between generations 7 and 31 (Fig. 1A). Therefore, following an initial rapid increase in 7 generations, the fecundity of the evolved populations had stabilized until generation 31. At the gene expression level, a considerable number of DEGs were shared between temporally separated evolved populations at generations 7 and 31 (Fig. 2). Furthermore, the magnitude and direction of the gene expression changes in the at generation 7 were highly correlated with those at generation 31 (Pearson correlation coefficient of log_2_FC = 0.96). These results suggest that the changes in molecular phenotypes have remained highly stable in evolved populations assayed 24 generations apart. The temporal patterns of gene expression (Fig. 2) are consistent with the patterns of fitness change (Fig. 1A), suggesting that after a rapid and pronounced change, the phenotype of populations stabilized after generation 7.

Although the selected traits are not known, we expect that the expression pattern reflects the dynamics of the selected traits. After a rapid response during the first generations, the expression of genes will stabilize and reach a plateau. To identify the genes matching this predicted expression pattern, we first identified DEGs by contrasting gene expression in the founder population with that in evolved populations in generations 7 and 31 (Fig. 2). Next, we classified the DEGs into 5 groups according to their conserved co-expression patterns. The DEGs shared between generations 7 and 31 were classified into 3 groups: ‘plateau’, ‘monotonic’, and ‘incomplete-reverse’ (Fig. 3, S8-9). Additionally, DEGs identified only in generation 7 or 31 were classified as ‘complete-reverse’ and ‘late-response’, respectively (Fig. 2-3, S8-9).

**Fig. 3.**
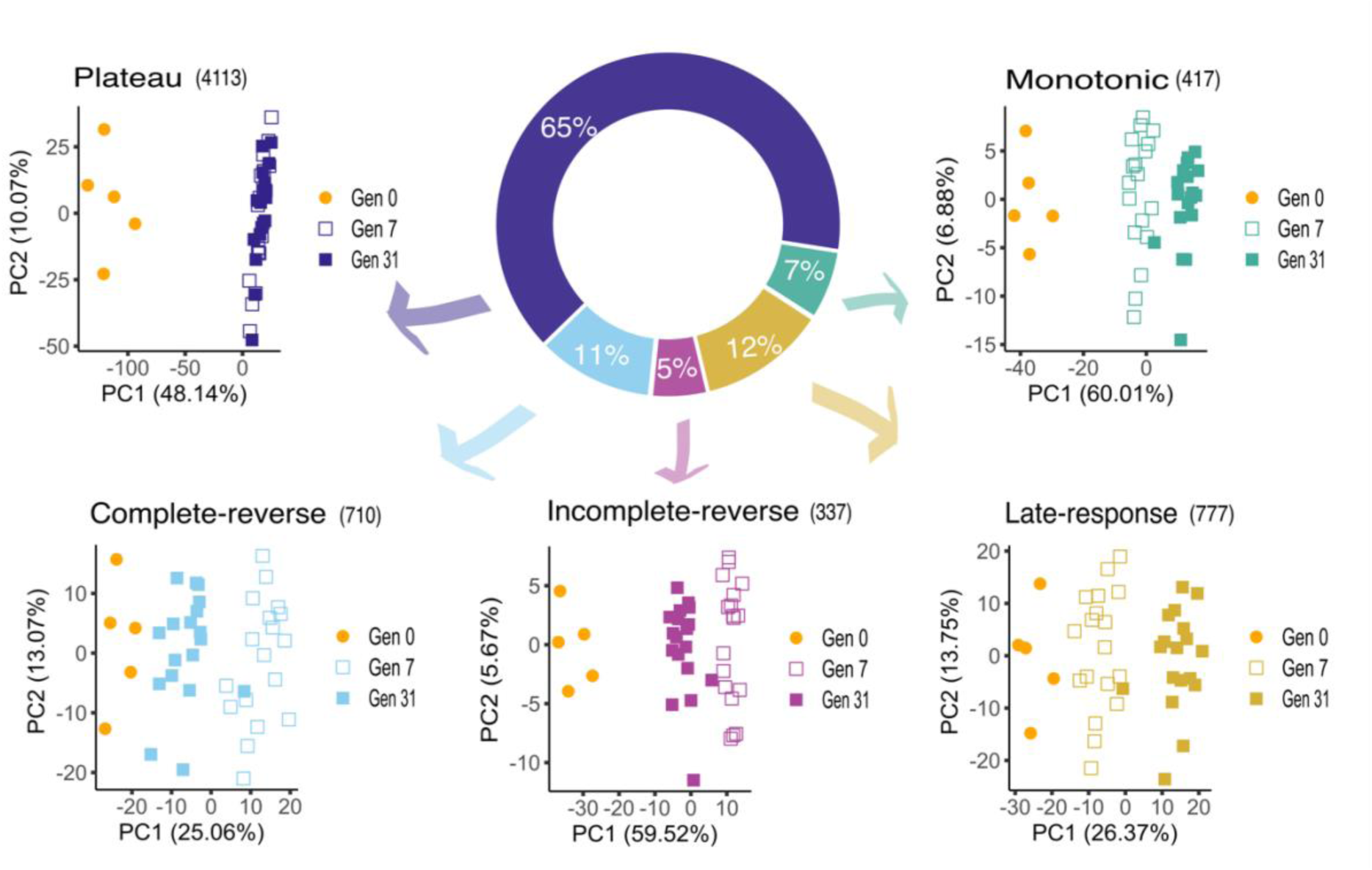
Evolutionary dynamics of differentially expressed genes (DEGs) with conserved co-expression patterns. The pie chart which illustrates the proportion of DEGs in each group highlights that the expression of the majority of genes (65%) have reached a plateau. Principal Component Analysis (PCA) shows that the expression of the ‘plateau’ group has stabilized at generations 7 as PC1 and PC2 cannot distinguish between the evolved populations at generations 7 and 31. PCA depicts the directional continuous change in the expression of ‘monotonic’ and ‘late-response’ groups throughout time. However, the magnitude of expression changes at generation 7 is larger in the ‘monotonic’ group than the ‘late-response’ one. For ‘complete-reverse’ and ‘incomplete-reverse’ groups, the direction of expression change reverts to that of the founder population. However, the magnitude of expression change from generation 7 to 31 is higher in the ‘complete-reverse’ group. PCA was conducted on the scaled log_2_ per million (log_2_CPM) values of the DEGs in each group separately. The number of DEGs in each group is indicated in parentheses. Gen0, Gen7, and Gen31 correspond to gene expression at generations 0, 7, and 31, respectively. Each point represents a sub-replicate sample.

We define the ‘plateau’ group as those genes that, after a significant expression change at generation 7, do not show significant change in expression between generations 7 and 31. The majority of DEGs (4,113 genes, 64.73%) belonged to the ‘plateau’ group (Fig. 3, S8-9). The large number of genes with plateaued expression supports the high correlation between the transcriptomic profiles in the evolved populations that were phenotyped 24 generations apart (Fig. 2, S4B). A total of 417 DEGs (6.56%) showed a monotonous change in their expression until generation 31 (‘monotonic’ group, Fig. 3); the expression change (either increase or decrease) of this group in the evolved populations was in the same direction in generations 7 and 31, and significantly different from that in the founder population. The ‘incomplete-reverse’ group (337 genes, 5.30%) exhibited a reversal in the direction of expression change after generation 7 toward the founder population (Fig. 3, S8-9); although the gene expression at generation 31 remained significantly different from that of the founder population. Identification of the other two groups of DEGs (‘complete-reverse’ and ‘late-response’) provides interesting insights into the transient gene regulatory responses during the early directional phase of adaptation. A total of 710 DEGs, (11.17%), classified as ‘complete-reverse’ group, were initially up (or down) regulated in the evolved populations after 7 generations, but reverted to the expression level that was not significantly different from the founder population at generation 31 (Fig. 3, S8-9). Additionally, the expression changes of 777 genes (12.2%), ‘late-response’ group, were in the same direction at generations 7 and 31 compared to the founder population, but a significant change was only identified at generation 31 (Fig. 2-3, S8-9).

The expression pattern of the ‘monotonic’ and ‘complete-reverse’ groups resembles that of the ‘late-response’ and ‘incomplete-reverse’ groups, respectively (Fig. S8). Despite the slight similarity of the expression trajectories of these groups, the PCA showed different patterns of differentiation between the founder and evolved populations (Fig. 3, S9). Additionally, the ‘monotonic’ and ‘late-response’ groups, as well as the ‘incomplete-reverse’ and ‘complete-reverse’ groups, showed significant differences in expression variance (Fig. S10, Table S3) and pleiotropy at the levels of gene connectivity and tissue specificity (Fig. 6, S15). The proportion of these five groups remained consistent even when only genes with absolute log_2_FC > 0.32 were considered, i.e. genes with an absolute change of at least 25% (Fig S11). This finding indicates that the predominant pattern was not driven by genes with small expression change.

Functional analysis revealed a large number of GO terms with diverse biological functions in each group (Table S2A, Fig. S5-S7). Genes in the ‘plateau’ group were enriched for GO terms such as ‘sexual reproduction’ (GO:0019953), ‘cytoplasmic translation’ (GO:0002181) and ‘renal system process’ (GO:0003014) (Table S2C, Fig. S5). In addition, the expression of the ‘plateau’ genes was enriched in rectal pads and testis (Table S2C, Fig. S7). Female fecundity increased significantly after 7 generations, followed by a plateau (Fig. 1A). The enrichment of ‘plateau’ genes for reproduction-related GO categories and male reproductive tissue, i.e. testis, matches the stabilization of female fecundity in the evolved populations (Fig. 1A).

## Adaptation through diverse patterns of co-expression in differentially expressed genes

Although the ‘plateau’ group comprises the most genes with expression changes, other groups of DEGs with conserved co-expression patterns also contribute to adaptation. The functional analysis of the monotonic genes showed enrichment of GO terms related to lipid metabolism such as ‘lipid catabolic process’ (GO:0016042), ‘fatty acid catabolic process’ (GO:0009062) and ‘fatty acid beta-oxidation using acyl-CoA oxidase’ (GO:0033540) (Table S2A, Fig. S5). To test whether the lipid content in the evolved populations followed the changes in expression of monotonic genes, we measured total lipid content in the founder and evolved populations from the same generations as the transcriptomic analysis. The lipid content in males in the evolved populations, similar to the expression pattern of the ‘monotonic’ group, gradually decreased compared to the founder population (Fig. 1C). The trade-off between fat storage and reproductive investment is demonstrated across various species, including both mammals and insects (16). Given the rapid and pronounced increase in fecundity (Fig. 1A), we expected to observe a reduction in the fat storage. The decrease in lipid content in females demonstrated a comparable pattern to males, however the lipid content in the evolved populations was not statistically significant from the founder population (Fig. 1B). The significant increase in the female fecundity and decrease in the lipid content (Fig. 1) matched the widely observed relationship between reproduction investment and fat storage.

Unlike other groups, the expression trajectory of the ‘incomplete-reverse’ and ‘complete-reverse’ groups was altered after the initial phase of adaptation with expression in generation 31 resembling that of the founder population (Fig. 3, S8-9). This pattern may be indicative of adaptation in response to the selection pressures experienced right after the environmental change. If the fitness costs of genes in ‘incomplete-reverse’ and ‘complete-reverse’ groups are high, their contribution could be substituted by other genes after the initial phase of adaptation. Overall, the expression trajectories of these genes demonstrate the robustness of gene regulatory networks. Interestingly, the trajectories of these genes have a comparable pattern to the changes in weight in both females and males (Fig. S3).

## Temporal magnitude of phenotypic evolutionary change

We determined the magnitude of phenotypic changes from the onset of environmental shift across high-order and molecular phenotypes between different time points using Darwin’s numerator (17). Darwin quantifies the rate of change in a trait as the difference in the natural logarithm of population mean per million years. The time intervals between our phenotypic assays are different (7 generations between the founder and evolved population at generation 7 versus 24 generations between the evolved populations at generations 7 and 31). Since the phenotypic changes over longer time intervals is slower (17), instead of Darwin’s value, we present the absolute values of Darwin’s numerator, i.e. the difference between log-transformed phenotypic means at different time points.

The magnitude of phenotypic changes for the high-order traits in the first 7 generations was greater than changes in other *Drosophila* evolution experiments (12, 18) in which smaller populations evolved for 39 to 103 generations (Fig. 4A). The phenotypic changes in fecundity observed in our populations were comparable to those in another evolution study (19) with large populations (see ‘Outdoor cages’ in Fig. 4A). We also observed heterogeneity in the magnitude of phenotypic response across traits (Fig. 4A) as changes in fecundity were greater than changes in the weight and lipid content measured in females.

**Fig 4.**
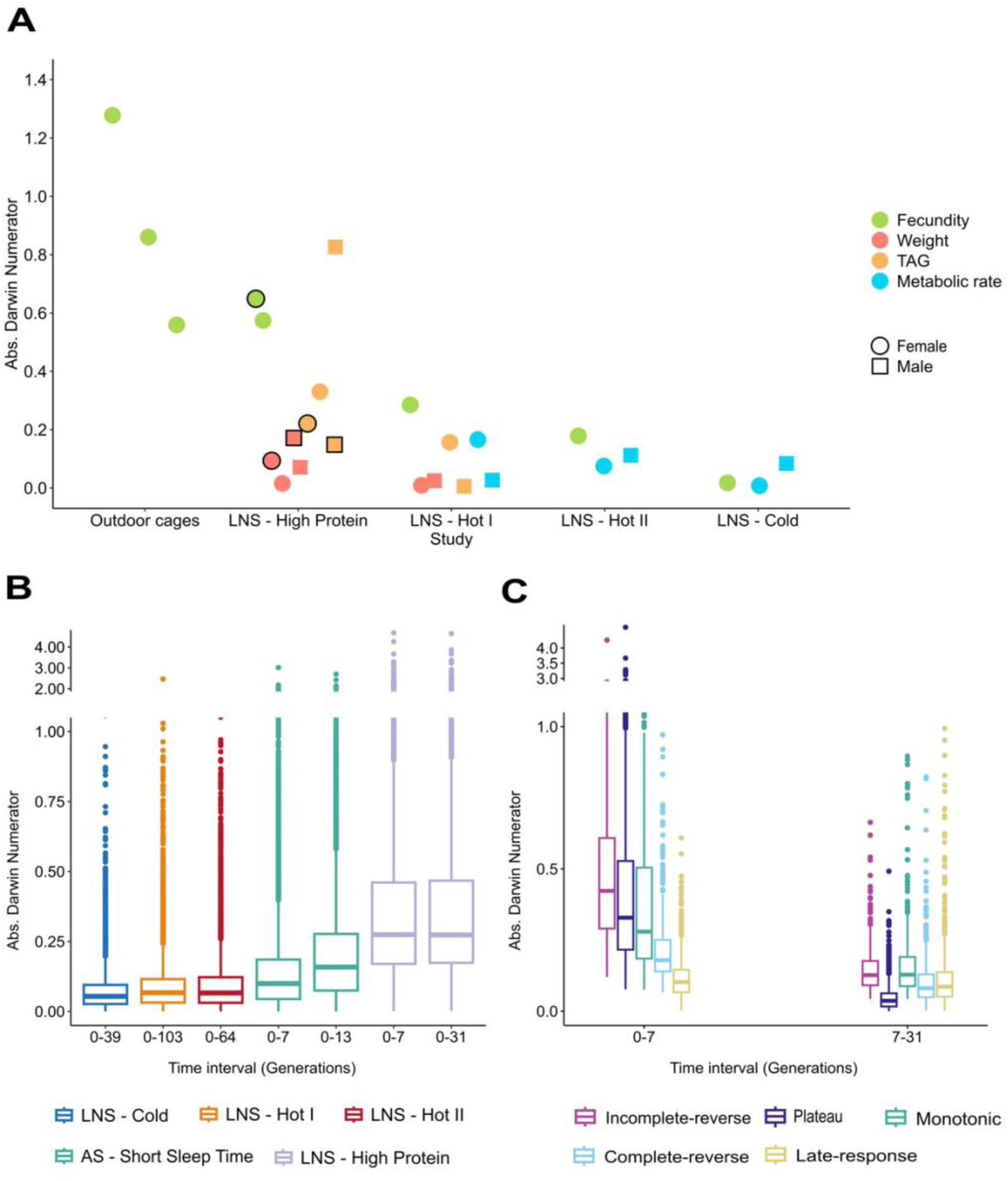
Rapid changes in fitness and gene expression after the environmental shift is followed by stasis. A) Magnitude of the phenotypic change for high-order phenotypes across evolution experiments in *Drosophila*. ‘Outdoor cages’: *D. melanogaster*, population size = 100,000, exposed to a fluctuating environment, and phenotyped at three time points spanning a total of ∼10 generations (19). Darwin numerators correspond to intervals of 3, 6, and 10 generations. ‘LNS - high protein’: the current study, *D. simulans*, population size = 100,000, adaptation on a high-protein diet for 31 generations. ‘LNS hot I’: *D. simulans*, population size = 1250, evolved in a new hot temperature regime for >100 generations (11). ‘LNS hot II’: *D. simulans*, population size = 1000, evolved in a new hot temperature regime for 64 generations. ‘LNS cold’: *D. simulans*, population size = 1000, evolved in a new cold temperature regime for 39 generations (12). In ‘LNS - high protein’, phenotypic changes between generations 0 and 7 are outlined in black. B) Magnitude of gene expression changes in *Drosophila* evolution experiments. ‘LNS Hot I’: the same population as in A, transcriptomic data from (18), ‘AS - Short Sleep Time’: *D. melanogaster*, population size = 25, truncation selection on sleep duration for 13 generations (20). C) The magnitude of gene expression change in groups with conserved co-expression patterns.

Transcriptomic changes in our study were more pronounced than in other evolution experiments, including those on the same species adapting to new temperature regimes over 30– 100 generations (12, 18) and a directional selection experiment (20) (Fig. 4B). While gene expression continued to shift under ongoing directional selection in Souto-Maior et al. (20) our populations showed rapid changes until generation 7, followed by a plateau through generation 31, indicating transcriptomic convergence (Fig. 4B).

Among the DEGs with conserved co-expression patterns, ‘incomplete-reverse’, ‘plateau’, and ‘monotonic’ groups had the highest phenotypic change in the first 7 generations followed by a reduction until generation 31 (Fig. 4C). The decrease in phenotypic change was especially pronounced for the ‘plateau’ group (Fig. 4C). Interestingly, ‘incomplete-reverse’ group showed the largest early phenotypic change (first 7 generations) before the shift in the direction of expression toward the founder population level (Fig. 4C). This suggests the ‘incomplete-reverse’ group had a transient role in adaptation and contributed to transcriptomic resilience. In contrast, ‘late-response’ genes showed the smallest phenotypic change in the early phase of adaptation (Fig. 4C), but the phenotypic changes remained consistent afterwards. Overall, the phenotypic changes were rapid after the environmental shift, followed by stagnation (Fig. 4).

## Highly parallel adaptive responses among replicate populations

As populations adapt towards a new optimal phenotype in an environment with a simple fitness landscape, we expect the replicates to reach phenotypic convergence near the new trait optimum. The levels of parallelism vary in population of different sizes (6, 19, 21), and across different biological levels e.g., genomic, transcriptomic and metabolomic (22). However, we expect that the degree of parallelism among replicates at the phenotypic level remains relatively stable when populations are close to the new optimum (6). We used temporal changes in parallelism among replicates as a proxy for proximity to the new optimum. If the replicates are near the optimum by generation 7 - as suggested by high-order (Fig. 1A) and molecular phenotypes (Fig. 2-3) - then parallelism should remain stable between generations 7 and 31.

For the high-order phenotypes, i.e. fecundity and weight, replicates converged after 7 generations (except 2 replicates), and there was no significant difference between six replicates at generation 31 (Fig. S12, Table S4). To compute the similarity, i.e. parallelism, of the DEGs between replicates, we first contrasted each replicate’s expression response to the founder population independently. Then, we computed similarity in DEGs between pairs of replicates using the Jaccard index, JI (23). The number of DEGs shared across all six replicates (2413 and 2399 genes at generation 7 and 31, respectively) are considerably higher than those that are specific to 1 to 5 replicates (Fig. 5A, S13). The average pairwise similarity between the replicates was relatively high at generations 7 (mean JI = 0.667 and 31 (mean JI = 0.666) (Fig. 5B). Moreover, parallelism among replicates did not differ significantly between generations 7 and 31 (two-tailed Wilcoxon Rank sum test, *p*-value = 0.68) suggesting that the replicates were already close to the new optimal phenotype as early as generation 7. To test whether the high degree of parallelism among replicates was driven by the number of DEGs, we generated a null distribution using delete-D jackknifing. The observed mean Jaccard index among replicates significantly exceeded the null expectation (Fig. S14), confirming that the parallelism was not affected by the number of DEGs.

**Fig. 5.**
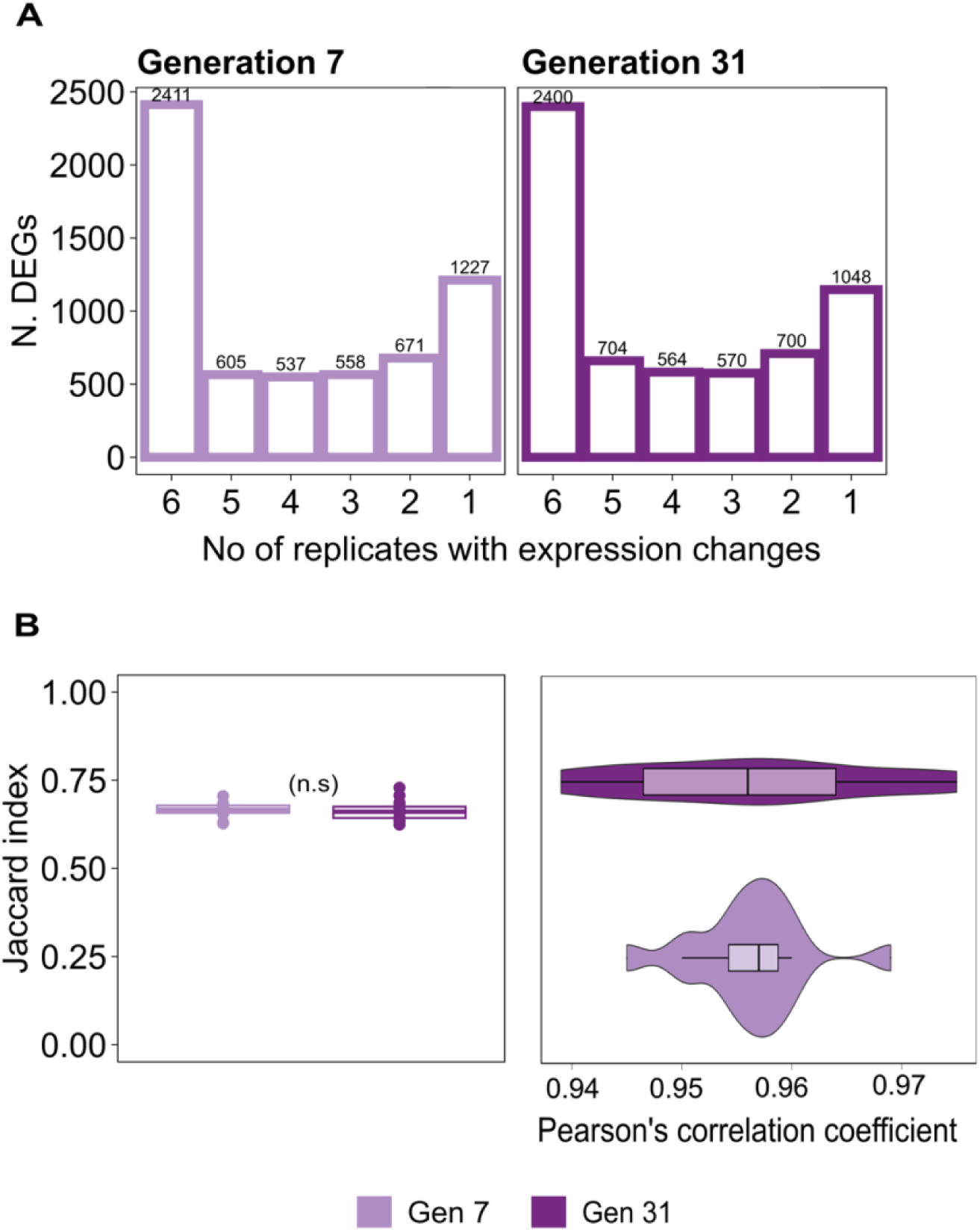
High similarity among the evolved replicate populations and the plateau of gene expression parallelism. A) The number of DEGs across replicates at generations 7 and 31 using replicate-specific analysis shows that the majority of identified DEGs are shared across six replicates. The x axis shows the number of replicates where a gene was identified as differentially expressed. B) Two metrics of similarity in the transcriptomic response: 1) Jaccard index (JI, left panel) shows the pairwise proportion of DEGs (JI = 0; no overlap, JI= 1; complete sharing). There is a high level of parallelism at both generations (mean JI = 0.66) with no significant difference between the two time points (Wilcoxson rank sum test, two-tailed, *p*-value = 0.68). 2) Pairwise Pearson’s correlation coefficients (Right panel) of gene expression response (log_2_FC) in evolved replicates at generations 7 (Gen 7) and generation 31 (Gen31).

As the identification of DEGs depends on the significance threshold of the differential expression tests, we additionally compared the magnitude and direction of gene expression change among the six replicates using Pearson’s coefficient of correlation (r). Transcriptomic responses were highly correlated between replicate pairs at each timepoint, with no significant difference between generations 7 and 31 (Fig. 5C), supporting strong phenotypic parallelism.

## Contribution of pleiotropic genes to different phases of adaptation

Pleiotropy is pervasive (24) but the extent to which it facilitates or constrains adaptation is debated. Some theoretical studies suggest that pleiotropy constrains adaptation due to the cost of complexity (25), while others suggests that, due to the high gene-trait modularity, pleiotropic genes can facilitate adaptation (26, 27). Here, to evaluate the extent to which pleiotropic genes contribute to phenotypic adaptation, we used two independent measures as proxies for pleiotropy for each gene: tissue specificity and level of connectivity in gene regulatory networks. Pleiotropic genes are less tissue-specific as they are involved in multiple functions in different tissues (24, 28). Pleiotropic genes also affect the expression of other genes thus have a higher number of connections in gene regulatory networks (29, 30). Using two measures of pleiotropy provided a complementary approach for estimating pleiotropy, enabling us to evaluate the robustness of the observed patterns to the proxy used. Genes that did not contribute to adaptation through change in expression (non-DEGs) were significantly more pleiotropic than genes with significant expression changes (Fig. 6A, S15, Table S5). There was also a negative correlation between the magnitude of change in gene expression and the degree of pleiotropy (both tissue specificity and gene connectivity) for all DEGs groups (Fig. S16, Table S6). However, parallel adaptation across all six replicates was driven by genes that were significantly less pleiotropic than those identified in 2 to 5 replicates only (Fig. 6C, S15C, Table S7).

**Fig. 6.**
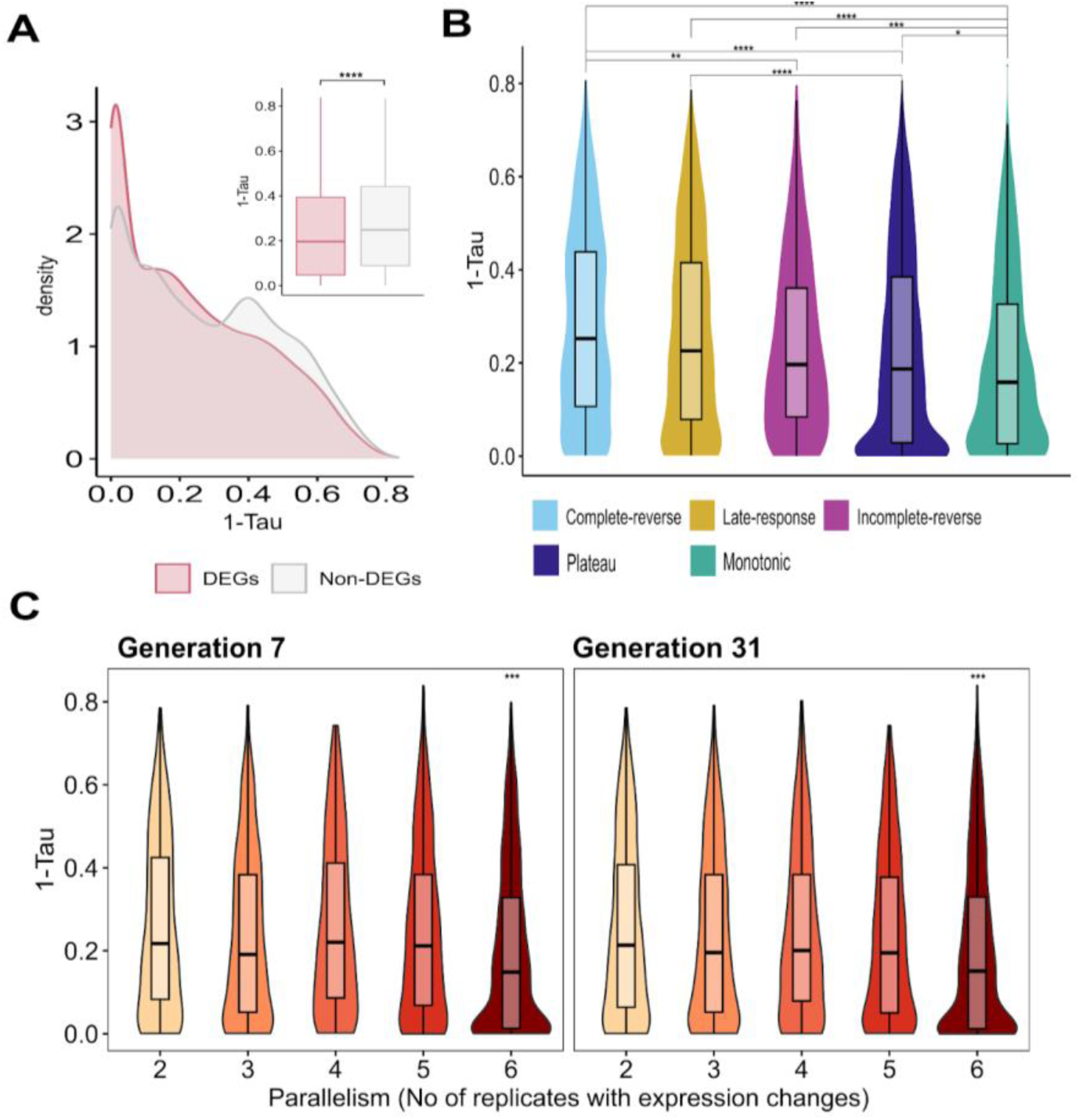
Low pleiotropic genes facilitate adaptation and drive parallel changes among replicate populations. A) Pleiotropy levels (tissue-specificity) of differentially expressed genes (DEGs) and non-differentially expressed genes (Non-DEGs) detected at generations 7 and/or 31 with available tissue specificity data (6,080 DEGs, and 3,682 non-DEGs). B) Pleiotropy levels of the DEGs groups with conserved patterns of co-expression. Two-tailed Wilcoxon rank-sum tests were conducted for two groups in (A) and all pairwise comparisons in (B). (*) *p*-value < 0.05, (**) *p*-value < 0.01, (***) *p*-value <0.001 and (****) *p*-value <0.0001. C) Pleiotropy levels (tissue-specificity) of DEGs across different numbers of replicates (e.g., group 2: DEGs shared by 2 replicates) show that genes with the highest level of parallelism (group 6) are significantly less pleiotropic than other groups (***, *p*-value <0.001).

The adaptive changes in fecundity (Fig. 1A) and gene expression (Fig. 2-3) that occurred by generation 7, followed by phenotypic stasis until generation 31, allowed us to conclude that the populations were close to the new optimum as early as generation 7. This prompted us to examine the contribution of pleiotropic genes to the different phases of adaptation. Genes in the ‘complete-reverse’ group, which contributed to adaptation only until generation 7 (Fig. 3) were significantly more pleiotropic than other groups using both proxies of pleiotropy (Fig. 6B, S15B). On the other hand, the ‘monotonic’ group where the change in expression level of genes did not reach a plateau even until generation 31, had significantly lower degree of pleiotropy (Fig. 6B, S15B). Apart from slight differences, tissue specificity and gene connectivity showed similar patterns (Fig. 6B, S15B). The contribution of ‘complete-reverse’ group with higher pleiotropy is transient and restricted to early phase of adaptation compared to genes with lower pleiotropy (‘monotonic’, incomplete-reversed’, and ‘plateau’ groups) which maintained their modified expression levels until generation 31.

## Discussion

Adaptation of polygenic traits is predicted to occur in two phases (4, 6, 7, 9); after a shift in the trait optimum, the mean phenotype changes rapidly towards the new trait optimum (directional phase). But the phenotypic change slows down to a pause when the population’s mean phenotype nears the optimum (stabilizing phase). Empirical studies to support these predictions require time-series phenotypic data to trace the changes in population’s phenotypic mean and keep the environment stable after the initial environmental shift. Additionally, replicates evolving in the same environment are needed to validate the repeatability of the phenotypic changes. Our evolution experiment was designed to meet these criteria. We measured the phenotypic states in six *D. simulans* replicate populations (100,000 individuals each) before and after an environmental shift. Using high-order and molecular phenotypic time series data, we confirmed the phenotypic patterns consistent with the predictions of polygenic adaptation. The directional phase of adaptation was characterized by fast phenotypic changes (Fig. 4). This was followed by the stabilizing phase, during which the magnitude of phenotypic changes in fecundity and gene expression decreased, indicating that the populations approached their new optimum (Fig. 4).

The phenotypic plateauing of the evolving populations is supported by several lines of evidence: 1) the adaptive fecundity, i.e. fitness, increase reaches a plateau after 7 generations (Fig. 1A), 2) a substantial fraction of DEGs (∼4,000 genes, 65%) have reached stable expression levels at generation seven, 3) the mean magnitude of gene expression change (measured as the Darwin’s numerator, Fig. 4B) remains constant between generations 7 and 31, and 4) the degree of parallelism among replicates has stabilized (Fig. 5, S12). The large proportion of ‘plateau’ genes suggests the proximity of the evolving populations to the new trait optimum, and hence a reduction in the strength of selection. In a truncation selection experiment where selection is constant throughout the experiment, the fraction of ‘plateau’ genes is expected to be smaller. Classification of the DEGs in a truncation selection experiment in *D. melanogaster* (20) based on their patterns of co-expression showed that the phenotypic changes were underlain by a large number of ‘late-response’ genes (55-61%), while the fraction of ‘plateau’ genes was rather small (12-15%) (Fig S17). The different proportion of the ‘plateau’ and ‘late-response’ genes in these two experiments supports our hypothesis that the populations may be at different distances to their optimal phenotypes. We note that while we detected significant changes in fecundity and gene expression at generation 7, these changes may have occurred earlier. In contrast, the lack of stabilization of expression changes for ‘monotonic’ and ‘late-response’ genes may suggest that these genes contribute to the traits for which their optimal values have not yet been reached, e.g. lipid content (Fig. 1B). Furthermore, these genes may play a role in fine-tuning gene expression and the phenotypic state of populations that are close to, but not quite at, the new phenotypic optimum.

The magnitude of evolutionary changes varied among different high-order traits e.g. fecundity, weight, and lipid content (Fig. 4). This may be due to trait-specific biological or physiological constraints, or to the different distances to their respective optima. The change in fecundity was greater than other high-order female phenotypes, supporting the idea that fitness components are highly evolvable (31). The value of a quantitative trait in a population at the fitness peak may represent the optimal phenotype in the environment (i.e. it may not be the maximum achievable value) or be constrained due to trade-offs with other phenotypes. We suspect that the observed stasis in fecundity (Fig. 1A) is less likely to be due to constraints. Firstly, the maximum fecundity measured in our study (18-20 eggs/female/day, Fig. S18) is lower than the reported fecundity in *Drosophila* (32). Secondly, the fecundity of one replicate at generation 7 (Fig. S12) was significantly higher than the other replicates. These replicates then converged at generation 31 due to a continued increase in fecundity.

The highly parallel phenotypic response among the replicates (Fig. 5) may have been due to the large population size of 100,000 individuals in each replicate, as well as the experimental set-up, e.g. round-robin crossing of the founder population. The adaptive responses are expected to be more parallel in larger populations (6, 19, 21). Rapid adaptation has been observed previously in large experimental populations (19). The round-robin crossing scheme used to generate the founder population may have resulted in more homogenous fitness values among individuals because it maximizes the maintenance of genetic variation by crossing the founder lines (33). Altogether, using large population sizes has improved our ability to infer the phenotypic patterns of polygenic adaptation in short evolutionary timescales as has been demonstrated by simulations (6).

Our findings support the idea that the contribution of pleiotropic genes to adaptation depends on the extent of their pleiotropy. Our results indicate that a high degree of pleiotropy can constrain adaptation. Genes that did not change in expression had significantly higher levels of pleiotropy than DEGs (Fig. 6A). This pattern mirrors findings from another experimental evolution study (34). Similarly, DEGs across multiple *Drosophilid* species tend to be more tissue-specific— i.e., less pleiotropic—than genes with conserved expression (35). Additionally, we showed that highly pleiotropic genes exhibited smaller changes in expression magnitude compared to less pleiotropic genes measured at the same time point (Fig. S16). This is consistent with a previous study, where gene expression changes in laboratory and natural *Drosophila* populations were negatively correlated with the level of pleiotropy (37). These findings support the constricting effect of pleiotropic genes due to the cost of complexity, as predicted by Orr’s model of adaptation (25). Building on Fisher’s geometric model (38), this model assumes universal pleiotropy and predicts that genes with stronger pleiotropic effects are less favoured by selection because their change may negatively impact many traits (25). On the other hand, our results support the idea that a certain degree of pleiotropy can facilitate adaptation. In our study, the major drivers of parallel adaptive phenotypic change were genes with low-to-intermediate pleiotropy (Fig. 6B, S15B). This finding is consistent with previous studies on grayling (36), sticklebacks (39), and *Drosophila* (42), which have shown that adaptive changes in gene expression were primarily driven by genes with low-to-intermediate pleiotropy. These findings align with predictions that the modularity in gene–trait networks can buffer the costs of pleiotropy, allowing pleiotropic genes to contribute to adaptation (26, 27).

The contribution of pleiotropic genes to adaptation also varied between short- and long-term responses. Genes with a higher degree of pleiotropy (‘complete-reverse’ group) compared to other groups contribute solely to the early phase of adaptation, when the population is far from the optimum. In contrast, genes with the least pleiotropic degrees (‘monotonic’ group) exhibit consistent expression changes, even during the stasis phase of adaptation (Fig. 6). Further empirical support for these pleiotropy-driven dynamics comes from a study (40) which found that the early adaptation is driven by highly pleiotropic genes. Pleiotropic genes affect multiple traits, and highly pleiotropic genes have a larger per-trait effect and a larger overall effect size (26, 27, 41). Orr’s model of adaptation predicts that adaptation of a population towards a new fixed optimum occurs in steps of decreasing sizes (42). Large effect alleles contribute to adaptation when the populations are far from the optimal phenotype. As populations approach the trait optimum, the contribution of small effect alleles increases. Orr’s model considers adaptation driven by successive fixations of de novo mutations. Recent theoretical work (4, 9) has studied the contribution of loci with varying effect sizes from standing genetic variation during polygenic adaptation, a scenario which matches our experiment. Hayward and Sella (4) demonstrated that the early directional phase was driven by moderate-to-large effect alleles, while the late stabilizing phase was shaped by moderate-to-small effect alleles. Thus, our findings are consistent with the theoretical model predictions (4, 9) that large-effect alleles primarily contribute to adaptation when the populations are far from the optimal phenotype and that the contribution of small-effect alleles increases as populations approach the new trait optimum.

Studies across diverse systems - including plants (43), sticklebacks (39) and *Drosophila* (34)-have shown that parallel evolution is driven by genes with high levels of pleiotropy. However, in other studies (44), similar to our findings, genes with low levels of pleiotropy have been repeatedly selected in replicate populations. These contrasting patterns are attributed to differences in the scale of adaptation: parallel evolution through highly pleiotropic genes is linked to local adaptation, while repeatability due to genes with low pleiotropy is attributed to global adaptation (43). However, other factors likely influence the relationship between pleiotropy and parallelism in rapid adaptation as in our study. The repeatability of selection for highly pleiotropic genes in independent replicates may depend on their synergistic or antagonistic effects. Highly pleiotropic genes with synergistic effects on multiple traits can facilitate repeatable adaptation across replicates. Whereas pleiotropic genes with antagonistic effects may constrain adaptation and hinder the adaptive response across replicate populations. If such trade-offs significantly affect fitness, selection may favour genes with fewer constraints, and parallel adaptation would be driven by low-to-intermediate level pleiotropic genes. A second factor involves the magnitude of shift in trait optima across traits. Typically, several traits are under selection during adaptation. When the selected traits have high covariance and therefore undergo similar shifts in optimum, highly pleiotropic genes with synergistic effect on selected traits may contribute more effectively to adaptation and be repeatedly favoured. However, if the magnitude or direction of the shift in optima differs among traits, the resulting trade-offs can reduce the likelihood that the same highly pleiotropic loci will be selected for across replicates.

## Materials and Methods

### Founder population

The outcrossed founder population was set up from 93 inbred lines from a North American *Drosophila simulans* panel (45) following a two-way round-robin crossing scheme (Fig. S1). A total of 93 crosses were set up from all inbred lines. Each cross was maintained in a bottle with roughly 100 individuals and were let to randomly mate for 15 generations to facilitate recombination and generation of new haplotypes. The crosses were maintained on a standard high-carbohydrate diet with a protein to carbohydrate (P:C) ratio of 1:4 (Fig. S2, Table S1) at 25°C, and a light cycle of 12:12 hrs light and dark. A total of 1075 individuals from each cross (total of 93 crosses) were pooled to establish the founder population with 100,000 individuals.

### Evolution experiment

The founder population was exposed to a novel high-protein diet with protein to carbohydrate (P:C) ratio of 1:1 (Fig. S2, Table S1). Six replicates were established from the founder population through sequential egg-laying periods as follows; the founder population with 100,000 individuals were allowed to lay eggs on 30 trays (3.6x15x 22cm depth, WEBER®) of high-protein (1:1 P:C) food for 2 days. After 2 days, the trays of food were used to set up one replicate. After establishing 3 replicates, the egg-laying period was extended to 3 days due to a decrease in fecundity as individuals aged. A total of 6 replicates were set up from the founder population. Each replicate was maintained with a census size of 100,000 individuals in a large mesh cage (120x60x60 cm, BugDorm-6M620 Insect Rearing Cage, Bugdorm) at 25°C, with a light cycle of 12:12 hrs light and dark, with non-overlapping generations. The cages were maintained in a room with controlled temperature and humidity (Fig. S19).

### Times-series phenotypic assays

#### Common garden experiments

We performed several phenotypic assays on 1) the founder population before exposure to the high-protein diet at generation 0 (gen0), 2) the replicates after 7 (gen7) and 31 generations (gen31) of adaptation. For the common garden experiment at generation 0 (CGE-0), equal numbers (172 individuals) of 93 round-robin cross were pooled (a total of 16,000 individuals) on standard high-carbohydrate diet, and the eclosed adults were used to set up 10 replicates (each roughly 1,600 individuals). To phenotype the evolving replicates at generations 7 (CGE-7) and 31 (CGE-31), around 1000 individuals were collected from each replicate, and used to set up 3 sub-replicates. At each time point (generations 0, 7, and 31) the populations were maintained in a common garden environment for two generations on high-protein (1:1 P:C) diet at low density (400 eggs per bottle) before phenotyping to minimize the transgenerational effects. Eclosed individuals after two generations of maintenance in a common garden experiment were allowed to mate for 24 hours. In the following 3 days, females and males were separated while anesthetized with CO_2_. After 2 days of recovery from CO_2_, 6-days-old flies were flash frozen in liquid nitrogen and stored at - 80°C. These samples were used for RNA sequencing and lipid content quantification.

#### Fecundity assay and statistical analysis

Freshly eclosed individuals after 2 generations of maintenance in common garden were collected to measure female fecundity. In CGE-0, we measured fecundity in 10 replicates of the founder population (each with 3 sub-replicates), thus a total of 30 sub-replicates. In CGE-7 and CGE-31, fecundity was measured in 3 sub-replicates for each of the 6 evolved populations, a total of 18 sub-replicates. For each sub-replicate, roughly 100 freshly eclosed individuals were transferred to an embryo collection cage (Cat. #59–101, Genesee Scientific, San Diego, California) with a Petri-dish containing black media. The Petri-dish was changed twice a day for a week, photographed and the number of eggs were counted following the protocol in (46). Fecundity was measured for 7 days which spans the age of individuals during the maintenance of the populations in the evolution experiment. Females and males in each sub-replicate were counted, dried at 60°C for 24 hours, and weighed. Fecundity was measured as the total number of eggs laid over seven days per female. Weight was measured as the average dry mass (mg) per fly.

To assess the significance of changes in *fecundity* between the founder and evolved populations we used a linear mixed model using R package lme4 (version = 1.1.35.5) with *generation* as a fixed categorical effect with three levels (0, 7, and 31). We included *replicate* as random intercept with six levels (replicates 1 to 6) and uncorrelated, dummy coded centered *generation* as random slope to account for the hierarchical covariance structure in our data. *Weight* was used as a fixed continuous effect.

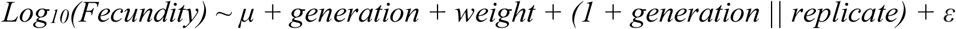

We also tested the divergence of *fecundity* among the 6 evolved replicates at each generation by fitting a linear model with *replicate* used as a fixed effect (with 6 levels). *Weight* was used as a fixed continuous effect. We fitted separate models for generation 7 and 31. We used the option REML = FALSE to request maximum likelihood estimation, which provides more accurate estimates for the fixed effects when comparing models.

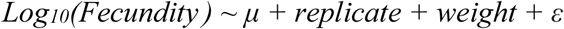

*fecundity* was log_10_-transformed to meet the assumptions of normality of residuals, and heterogeneity of variance. To test for the significance, we performed pairwise post-hoc tests using function *emmeans* from package *emmeans* with option *pairwise*, and corrected the *p* values applying the Tukey’s honest significant difference (HSD).

We also tested for the significance of changes in *weight* between the founder and evolved populations using a linear mixed-effects model with option REML = FALSE. We included *generation* as a fixed categorical effect with three levels (0, 7, and 31), and *sex* as a fixed categorical effect with two levels (female, male), along with their interaction. To make the variance homogeneous, we aggregated sub-replicate data by computing the mean weight for each replicate, sex, and generation combination. We included r*eplicate* as a random intercept with six levels (replicates 1 to 6). We used the to request maximum likelihood estimation, which provides more accurate estimates for the fixed effects when comparing models.

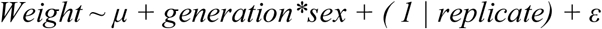

To test the divergence of *weight* among the replicates at each generation we fitted a mixed-effects model with *replicate* as a fixed effect with six levels (replicates 1 to 6). We fitted separate models for generations 7 and 31, and for each sex independently because the full model did not meet the assumption of homogeneity of variance. *weight* was log_10_-transformed.

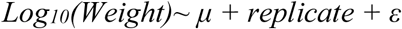

To test for the significant phenotypic changes in *weight*, pairwise post-hoc tests was performed as described above. *P*-values across all contrasts were corrected using FDR (47).

### Lipid content assay and statistical analysis

We measured the amount of triglyceride (TAG) as the main constituent of body fat using coupled colorimetric assay as described in (11). Lipid content was measured in 8-10 sub-replicates for each sex of the founder population in CGE-0. For the evolved populations, lipid content was measured in 3 sub-replicates of each replicate for each sex in CGE-7 and CGE-31. For each sub-replicate, a total of six to nine 6-days-old flies were weighed and homogenized in 500µl PBST (1% Triton X-100) using an electric mortar (VWR®). The homogenates were incubated for 10 minutes at 70°C to inactivate enzymes, then centrifuged at 350 rpm for 3 minutes at 4°C. The supernatant was frozen at -80°C until quantification. We used a glycerol standard (Sigma G7793) with the following concentrations (1.2, 1, 0.8, 0.6, 0.4, 0.2, 0.1, 0 mg/mL) as reference. A total of 50 μL of supernatant and standard was transferred to a well in a 96-well plate. Samples were measured in duplicates. The initial absorbance was measured at 540 nm with the EnSpire 2300 Microplate Reader (PerkinElmer). Then, 200 μL of Triglyceride Working Reagent (TR22421 Thermo Fisher) was added to each well. The mixture was incubated at 37°C for 30 minutes with gentle agitation. The final absorbance was measured at 540 nm. All measurements were blank-corrected. The difference between the initial and final absorbance was attributed to the change in absorbance resulting from the conversion of triglyceride. Standard curves were made by fitting an inverse polynomial regression line, and were used to estimate the concentration of TAG. The technical replicates of each sample were used to calculate the coefficient of variation (CV) where *S1* and *S2* are the concentrations of technical replicates 1 and 2, and *m* is the mean. The concentration of TAG in samples with CV > 10% were repeated. The estimated amount of TAG for each sample was the average of its two technical replicates.

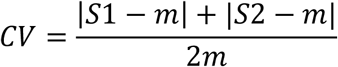

The lipid content was expressed as μg TAG equivalents per fly. We tested for the significance of changes in lipid content between the founder and evolved populations using a linear mixed-effects model with option REML = FALSE. We included *generation* as a fixed categorical effect with three levels (0, 7, and 31), and *sex* as a fixed categorical effect with two levels (female, male), along with their interaction. We used *weight* (wet mg per fly) as a fixed continuous effect. We aggregated sub-replicate data similar to above. *TAG* was log_10_-transformed. *Replicate* was included as a random intercept with six levels (replicates 1 to 6). The significance of phenotypic changes in TAG, and correction of *p*-values were done as described above.

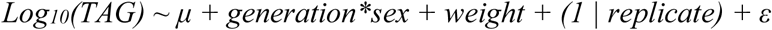

### Transcriptome analysis

#### RNA extraction and sequencing

We extracted total RNA from 5 sub-replicates of the founder population (CGE-0), 18 sub-replicates (3 sub-replicates for each replicates) in CGE-7 and 18 sub-replicates in CGE-31. For each sample, RNA was extracted from fifty 6-days-old males using TRIzol™ Reagent (Invitrogen) following the manufacturer protocol, and the reagent volume was adjusted to the sample. The extracted RNA was frozen at -80°C. RNA libraries were constructed using NEB poly-A enrichment multiplexed with dual indices. All samples were sequenced on the same lane on Illumina NovaSeq S4 PE150 XP platform.

#### RNA-Seq data processing

The paired-end 150bp reads were mapped to *D.simulans* reference genome (48) using GSNAP (version = 2023.03.24) (49) with parameters -A SAM -k 15 -N 1 -m 0.08. Read quality was checked for 3ʹ bias using RSeQC (Version 5.0.1) (50), the gene body coverage was similar in all samples. The number of reads aligned to the exons was counted using *featureCounts* function of *subread* (version = 2.0.3) (51) with parameters: -T 8 -s 0 -p --countReadPairs -B -t exon -g gene_id -Q 10 -C. The mapping statistics generated using SAMtools (52) with option -stats (version = 1.16.1), and the statistics from *subread* mapping are presented in Table S8-S9, respectively.

### Differential gene expression analysis

Gene expression data were normalized by library size using the *cpm* function in *edgeR* package (version 4.2.2) (53), and presented as counts per million (CPM). Genes with low expression were filtered out, keeping only those with at least one CPM in all samples. We used the *voomWithDreamWeights* function in the *variancePartition* package (version 1.23.3) (54) to compute log_2_-transformed CPM and estimate weights. We modeled the effect of evolution on gene expression, by fitting a linear mixed model (‘overall-model’) for each gene using *dream* function in *variancePartition.* We used log_2_ gene expression (*Y*) as the response variable, *generation* (0, 7 and 31) as a fixed categorical effect, and *ε* was the random error. Each replicate was phenotyped multiple times (at generation 7 in CGE-7 and at generation 31 in CGE-31), and a total of 3 sub-replicates were measured for each replicate. To account for this hierarchical covariance structure, we used an uncorrelated random intercept, random slope model. Random slopes were dummy coded and centered.

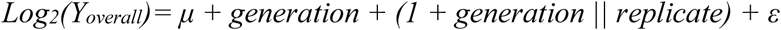

We used the *eBayes* function in the *variancePartition* package on the output of the *dream* function to compute moderated t-statistics and reduce the number of false positives. We performed differential gene expression analysis using three contrasts: a) generation 7 (CGE-7) vs. generation 0 (CGE-0), b) generation 31 (CGE-31) vs. generation 0 (CGE-0), and c) generation 31 (CGE-31) vs. generation 7 (CGE-7). The significance level was set at 5% after multiple testing correction using FDR (47) across all three contrasts. The magnitude of the gene expression change was expressed as the log_2_ fold change (log_2_FC).

Additionally, we assessed the replicate-specific gene expression response for each replicate separately (‘replicate-specific models’). We used log_2_ gene expression (*Y*) as the response variable, and *generation* as a categorical fixed effect with 3 levels (0, 7, and 31).

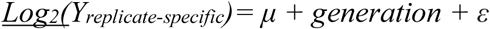

Similar to the ‘overall-model’ we performed differential gene expression analysis using three contrasts. Multiple testing corrections were performed across three contrasts and all 6 replicates. This ensured that the multiple testing correction to be as stringent as in the ‘overall-model’, i.e., with a similar number of tests. There was a high overlap in terms of the identified DEGs by the overall model and the replicate-specific models (Fig. S20A).

## Patterns of co-expression in differentially expressed genes

The DEGs were classified into five groups based on the significance and direction of change in expression as determined by the differential gene expression analysis model.

1. The ‘plateau’ group contained genes that their expressions significantly changed in both evolved time points, CGE-7 and CGE-31, compared to the founder population, CGE-0 (*p* value < 0.05), but no significant change was observed between two evolved time points, CGE-7 and CGE-31 (*p* value > 0.05).
2. Genes in the ‘monotonic’ group, similar to the ‘plateau’ genes showed a significant change in expression at both evolved time points, CGE-7 and CGE-31, compared to the founder population, CGE-0. But unlike the ‘plateau’ genes, the expression between CGE-7 and CGE-31 was significant. The direction of change in the expression of these genes (either decreasing or increasing) was consistent across all time intervals.
3. Genes in the ‘complete-reverse’ group had significant expression changes in CGE-7 compared to CGE-0. After CGE-7, the expression level reverted to the level observed in CGE-0.
4. The gene expression patterns in the ‘incomplete-reverse’ group were similar to the ‘complete-reverse’ group. However, the expression changes between the last time point (CGE-31) and the founder population (CGE-0) were significant (*p*-value < 0.05).
5. Genes in the ‘late-response’ group had significant expression changes in the last time point (CGE-31) compared to the founder population (CGE-0). However, the expression of these genes was not significant in the first time point (CGE-7) compared to the founder population (CGE-0). Similar to the ‘monotonic’ group, the direction of gene expression changes was consistent across all time intervals.

To ensure that the patterns of co-expression were not driven by genes with small changes in expression, we classified only the DEG with absolute expression change higher than 0.32 into groups using the above criteria. Additionally, we performed a similar classification of DEGs for the temporal male transcriptomic data in a truncation selection experiment in *Drosophila* (20). We first modelled the gene expression across 14 time points, then performed contrasts using time intervals that best aligned with our experiment: generation 0 to 7 (fully matching our early evolution phase), from generation 7 to 13 (the latest available time point) as well as the contrast between generations 7 and 13. The classification of DEGs into groups with conserved patterns of co-expression was done using the above criteria.

## Gene ontology, pathway, and tissue enrichment analyses

We used the topGO package (*version = 2.56.0*) (55) to perform gene ontology (GO) enrichment analysis. We ran a weighted Fisher’s exact test for each gene set using the *Weighted01* algorithm. We performed tissue enrichment using the expression data for 15 tissues in males available in the *FlyAtlas2* dataset (56). We first determined the tissue-specific genes by comparing the tissue gene expression levels with that of the whole-body, and genes with more than 2-fold expression in a certain tissue were classified as tissue-specific. Subsequently, we performed Fisher’s exact test for each tissue and gene set. Pathway enrichment was conducted with *clusterProfiler* package (version= 4.10.0) (57) using the Kyoto Encyclopedia of Genes and Genomes database (KEGG) (58) The background lists for all enrichment tests consisted of all expressed genes (10,197 genes). *P*-values were corrected using FDR (47) across all contrasts in tissue and pathways enrichment. We did not correct *p*-values from GO enrichment as suggested by the developers of topGO (55). **Magnitude of phenotypic evolution (Darwin numerator)**

We computed the magnitude of phenotypic changes in Darwin, which expresses the evolutionary change as a factor of the base of natural logarithm (*e*) per million years. The time intervals between our phenotypic assays are not equal (7 versus 24 generations), therefore, following (17) we calculate Darwin numerator (the phenotypic change between different timepoints in natural-logarithm scale). We present the absolute Darwin numerator as the magnitude, not the direction, of phenotypic change is relevant. We computed Darwin numerator using the following formula (17) where *y*2 and *y1* are the mean values of the log-transformed trait measured at different generations.

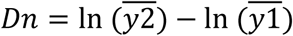

We quantified Darwin numerator for two time intervals: 1) between generation 0 and 7, and 2) between generation 0 and 31. To compute Darwin numerator for fecundity, weight and lipid content, we used the mean phenotype across six evolved replicates where the phenotype of each replicate is the average of 3 sub-replicates. The phenotype of the founder population is the average of 8-10 sub-replicates. For gene expression, Darwin numerator was computed for each gene separately. The gene expression in each evolved replicate was summarized as the mean expression level (CPM) across 3 sub-replicates. The expression of each gene at each time point is the mean expression level across six evolved replicates. Gene expression in the founder population is the average of gene expression in 5 sub-replicates. Note that all phenotypes were log-transformed before calculating the means.

## Parallelism in patterns of gene expression

We quantified the similarity of DEGs among the replicates, i.e. parallelism, using Jaccard index (23). We identified the DEGs independently in each replicate in generations 7 and 31 (CGE-7 and CGE-31) compared to the founder population (CGE-0). The Jaccard index was calculated using the following formula, where A and B are any two replicates:

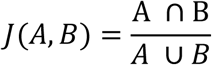

At each time point, all pairwise comparisons were performed between six evolved replicates. A two-sided Wilcoxon rank-sum test was used to assess the difference between generations 7 and 31. Additionally, we quantified the similarity between the replicates based on the correlation in their transcriptomic response. We computed Pearson’s correlation coefficients of the expression response (log_2_FC) using the full transcriptome after filtering lowly expressed genes for each replicate dataset (the numbers of genes after filtering ranged from 10397 genes to 10476 genes).

To ensure that the high parallelism between replicates was not due to the high number of DEGs, we generated 1000 sets each consisting of six replicates using delete-D jackknifing. In each set, the number of DEGs for each replicate matched the number of DEGs identified in ‘replicate-specific models’, which ranged from 3654 to 4283 genes, but was randomly drawn (without replacement) from the total expressed genes after filtering out low expressed genes (n = 10197 genes). In each set, the pairwise Jaccard index was computed between six evolved replicates. We compared the empirical mean Jaccard Index with this null distribution.

## Pleiotropy degree of differentially expressed genes

We quantified the degree of pleiotropy using two independent indices: network connectivity and tissue specificity. Genes with a higher number of connections in regulatory networks affect the expression of many gene, and are more pleiotropic (29, 30). In a gene regulatory network, the pleiotropic degree of a gene is determined by the number of genes that are affected by the focal gene. Thus, gene connectivity, i.e. the sum of all connections for each gene in a gene regulatory network, can be used as a proxy of pleiotropy (36, 37, 39). We computed the number of connections for each gene using the reconstructed gene regulatory network in *D.melanogaster* (59). Moreover, pleiotropic genes are expressed in more tissues as they are involved in multiple functions in different tissues (28, 60). Tissue specificity (tau) is a proxy of pleiotropy which measures the homogeneity of expression for a given gene across different tissues. We measured tissue specificity (tau) as described in (60) using tissue-specific gene expression dataset in FlyAtlas2 (56). We used male expression data from 15 tissues and excluded genes with zero expression across all tissues.

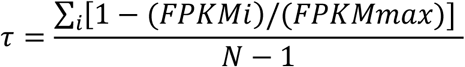

*τ* (Tau) is the coefficient of tissue specificity for a set of tissues (*N*), *FPKMi* (Fragments Per Kilobase of transcript per Million mapped reads) is the normalized expression value of a gene in tissue *i*, *FPKMmax* is the maximum expression value of the gene across the assessed tissues, and *N* is the number of assessed tissues. Tau ranges from 0 (uniform expression across tissues) to 1 (expression specific to one tissue). 1-Tau shows the degree of pleiotropy.

We measured pleiotropy using both indices for all DEGs and the five co-expression pattern groups. We also computed the Kendall’s tau rank correlation between the absolute expression change (log_2_FC) and pleiotropy for each transcriptomic group and generation. Furthermore, we tested whether the degree of pleiotropy is associated with the number of replicates in which a given gene is differentially expressed, i.e. the level of parallelism. We only included the DEGs that were identified in 2 replicates and had the same direction of expression change across the replicates. The significance of the degree of pleiotropy between different groups was tested using the Wilcoxon rank sum test, and *p*-values were corrected using FDR (47).

## Data analysis and visualization

All data analysis, statistical tests, and visualization were performed in R Statistical Software (v4.3.0, R Core Team).

## Data availability

The raw RNA-Seq reads will be available from the European Nucleotide Archive under project PRJEB83201 (accession numbers in Table S10) upon publication. Phenotypic data (fecundity, weight, lipid content, RNA count table) will be released upon publication. All scripts are available at https://github.com/ClaudiaRamL/BigCages_high_protein.

## Acknowledgement

We thank Paula Marconi and Magdalena Reitbauer for their help in maintaining the experimental populations, and Davide Salati for his help in the lipid measurements. We are grateful to Alexander Fedosov for his immense help in making many custom-made tools needed to maintain the large populations. We thank Claus Vogl and Andreas Futschik for their constructive suggestions, and all members of the Institute of Population Genetics for helpful discussion. RNA library preparation and sequencing were performed by the Next Generation Sequencing Facility at Vienna BioCenter Core Facilities (VBCF), member of the Vienna BioCenter (VBC), Austria. This research was funded by the Austrian Science Fund (FWF) to N. B. (P32672 and 10.55776/F91) and by the Max Planck Society. For open access purposes, the author has applied a CC BY public copyright license to any author-accepted manuscript version arising from this submission. The funders had no role in the study design, data collection and analysis, decision to publish, or preparation of the manuscript.

## Supplementary figures

**Fig. S1.**
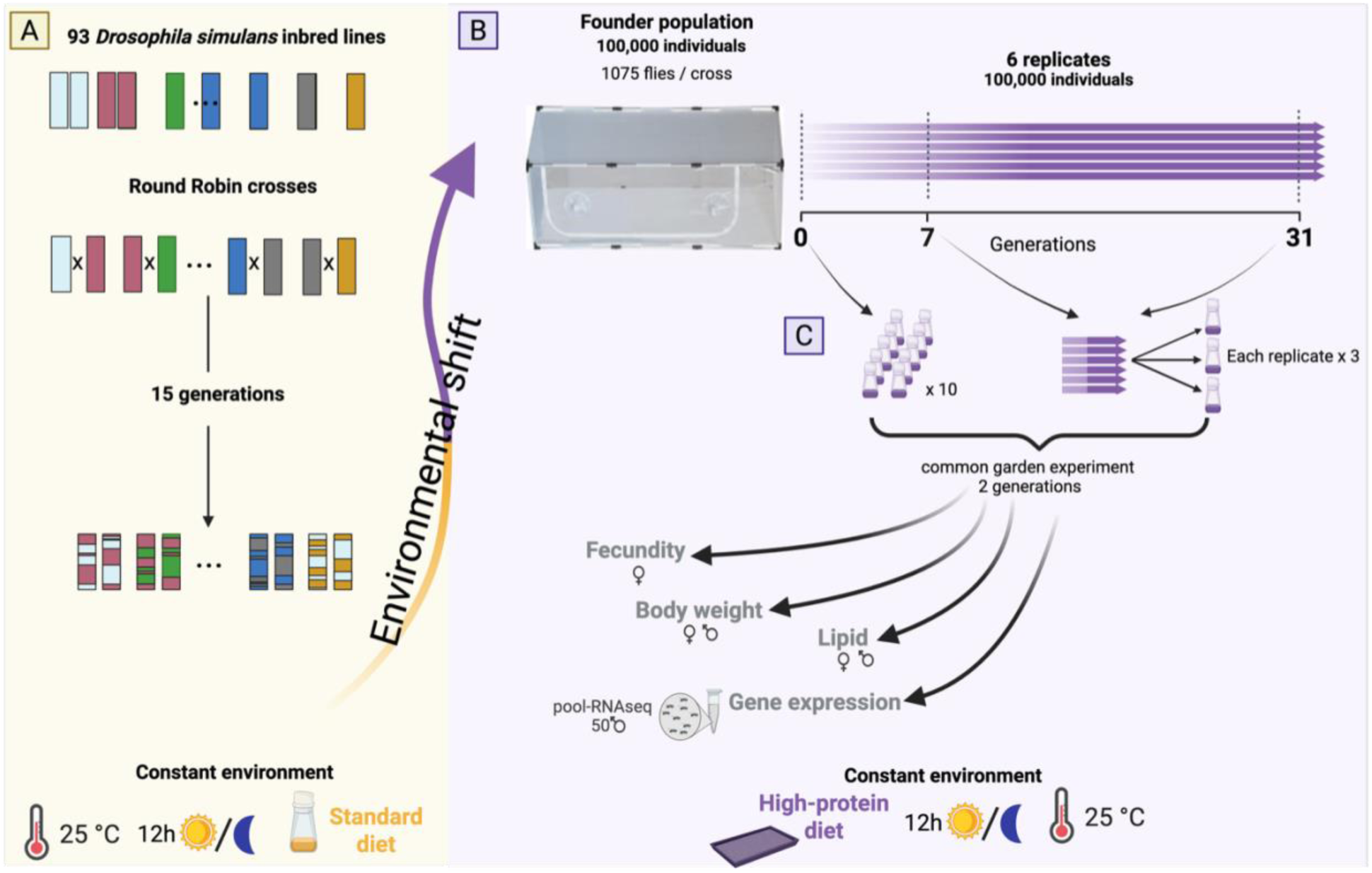
Experimental design. **A)** 93 inbred lines from a North American *Drosophila simulans* panel were used for round-robin crossing. 93 crosses were set up and maintained in small populations for 15 generations. The crosses were maintained on a standard high-carbohydrate diet with a protein to carbohydrate (P:C) ratio of 1:4 at 25°C, and a light cycle of 12:12 hrs light and dark. **B**) A total of 1075 individuals from each cross (total of 93 crosses) were pooled to establish the founder population. Six replicates were established from the founder population, each with 100,000 individuals. The replicates were maintained with a census size of 100,000 individuals in a novel high-protein diet with protein to carbohydrate (P:C) ratio of 1:1 for 40 non-overlapping generations. **C**) At three time point (generations 0, 7, and 31) the populations were phenotyped. The populations were maintained in a common garden environment on high-protein (1:1 P:C) diet for two generations before phenotyping. At generation 0, 10 replicates of the founder population were set up. To phenotype the evolving replicate populations at generations 7 and 31, around 1000 individuals were collected from each replicate population, and used to set up 3 sub-replicates. Eclosed individuals after two generations of maintenance in a common garden experiment were allowed to mate for 24 hours. 6-days-old flies were flash frozen in liquid nitrogen and stored at - 80°C. These samples were used for RNA sequencing and lipid content quantification. Freshly eclosed individuals after 2 generations of common garden were collected to measure female fecundity and body weight.

**Fig. S2.**
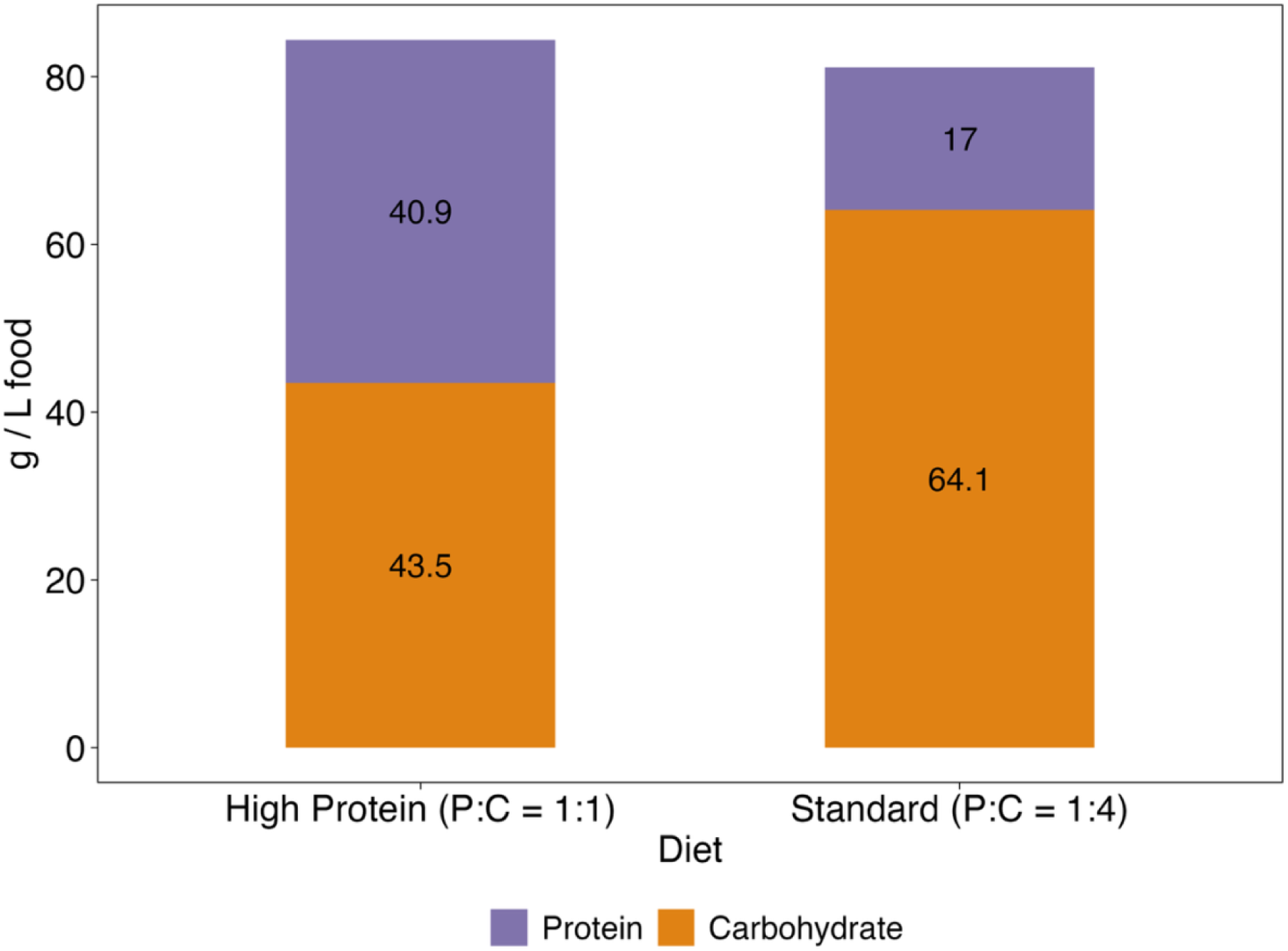
The amount of protein and carbohydrate in the standard (1:4 protein:carbohydrate P:C) and high-protein (1:1 P:C) diets. The high-protein diet with a protein:carbohydrate (P:C) ratio of 1:1 had 2.4 times more protein and almost 1.6 times less carbohydrate than the standard diet with a P:C ratio of 1:4. See Table S1 for the recipes and ingredients of the 1:4 and 1:1 P:C diets.

**Fig. S3.**
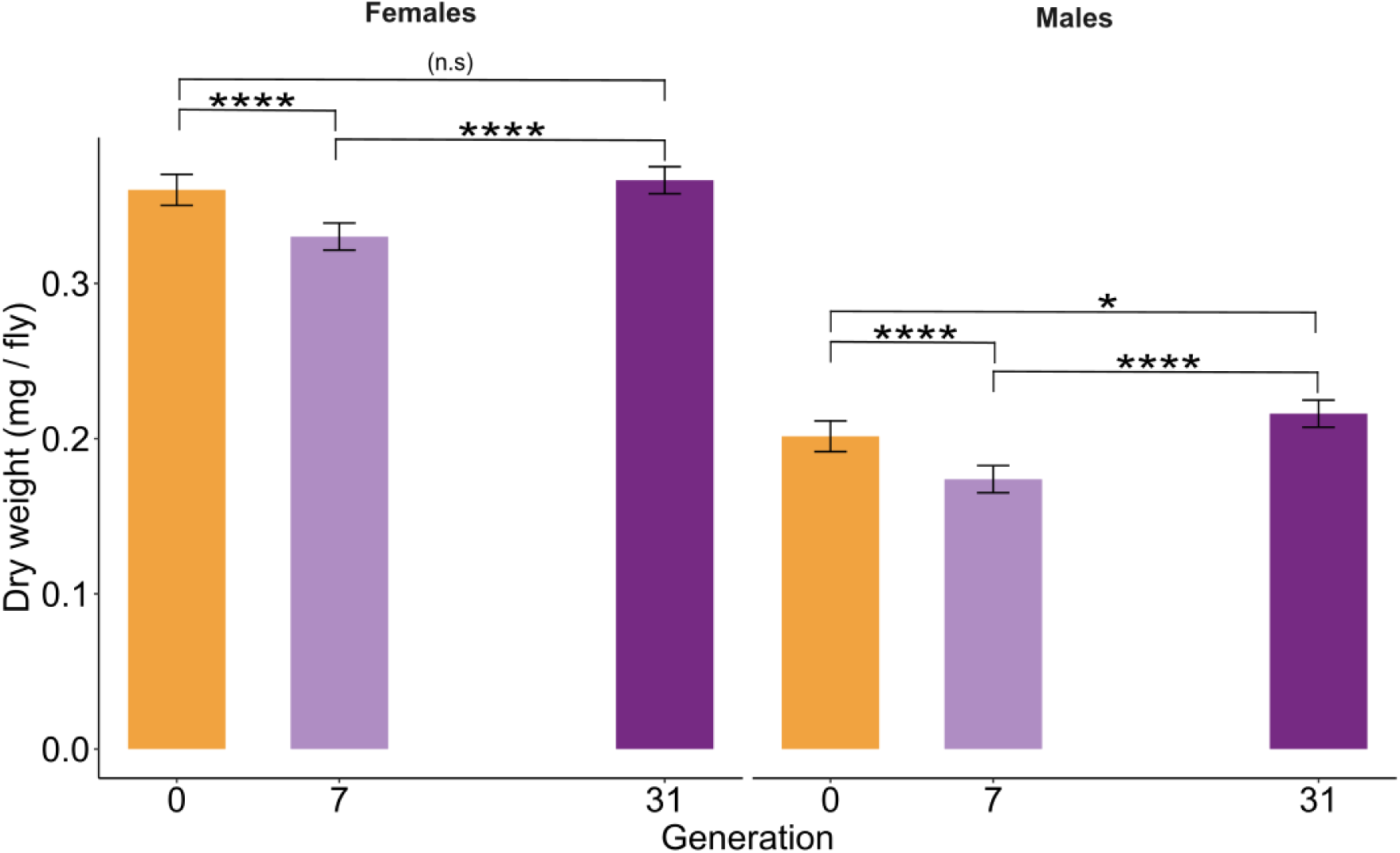
The adaptive changes in body weight. Body weight decreased significantly in both females and males at generation 7. Then, the direction of the change reversed in both sexes. Females increased in weight from generation 7 to generation 31, reaching the level of the founder population. Males also increased significantly in weight from generation 7 to 31, becoming heavier than the founder population. Significance was assessed using emmeans with option *pairwise* option. *P*-values were corrected across all contrasts using the Benjamini-Hochberg’s false discovery (FDR) method (Benjamini & Hochberg, 1995). Asterisks indicate significance levels: *p*-value < 0.05 (*), *p*-value < 0.01 (**), *p*-value < 0.001 (***), and *p*-value < 0.0001 (****). The bars depict the predicted means of the linear model and error bars show the 95% confidence intervals of the least-square means. See Table S4 for the result of all statistical tests.

**Fig. S4.**
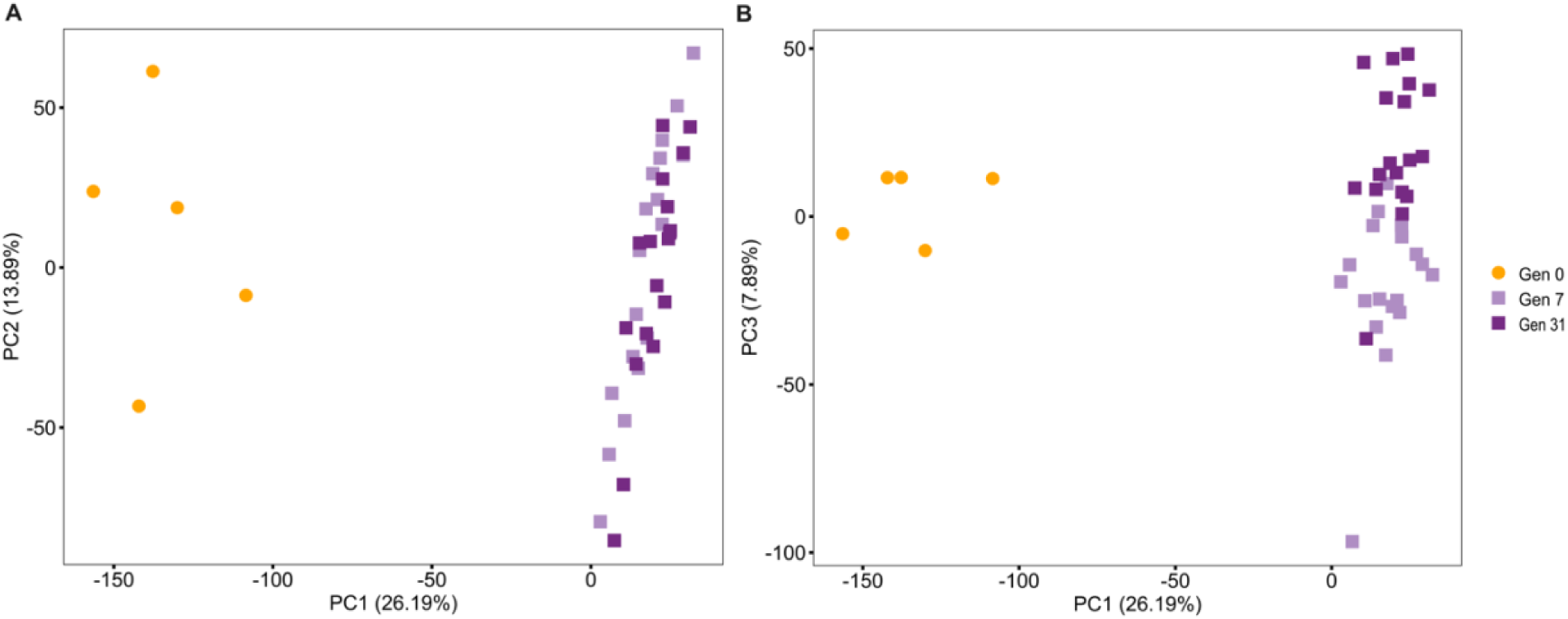
Transcriptomic divergence of the founder and evolved populations after 7 and 31 generations of adaptation. PC1 explains 26.19% of the variance and differentiates the founder population from the evolved populations (A). PC3 which accounts for 7.89% of the variation separates the evolved populations at generations 7 and 31 (B). Overall, the principal component analysis (PCA) shows a clear divergence between the founder and evolved populations, despite less differentiation between the evolved populations. PCA was performed using the normalized and scaled counts per million (CPM) of all genes after removal of the lowly expressed genes (10,197 genes). Gen 0: founder population, Gen 7: evolved replicate populations at generation 7, Gen 31: evolved replicate populations at generation 31.

**Fig. S5.**
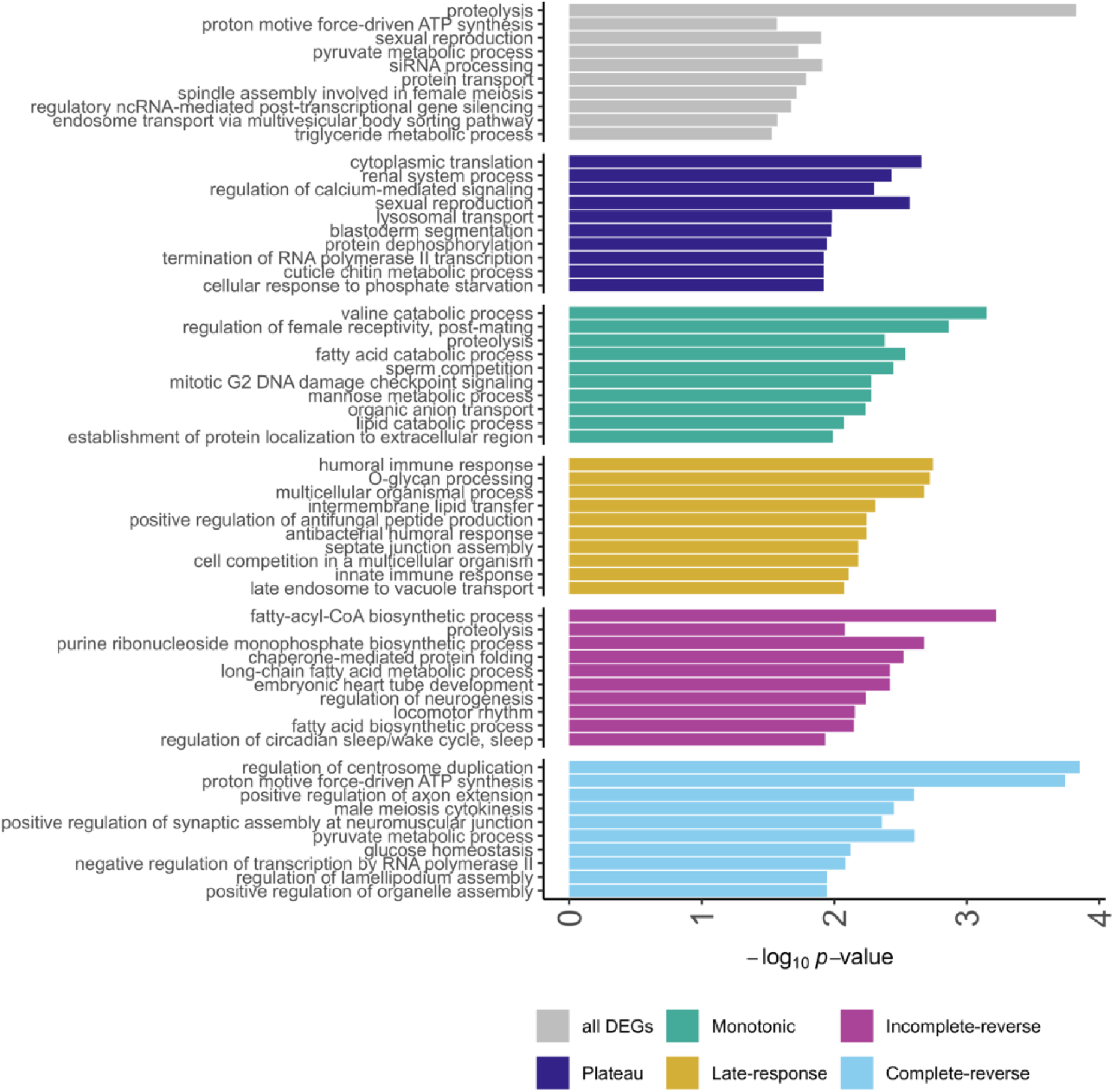
Enrichment of gene ontology (GO) terms across differentially expressed genes with conserved co-expression patterns. The top 10 most significant GO terms with the smallest *p*-values are shown for each group. The ‘all DEGs’ group refers to all the differentially expressed genes identified in generations 7 and/or 31. The -log_10_-transformed *p*-values of the enrichment test are shown on the x axis. For the complete list of GO terms and associated *p*-values see Table S2.

**Fig. S6.**
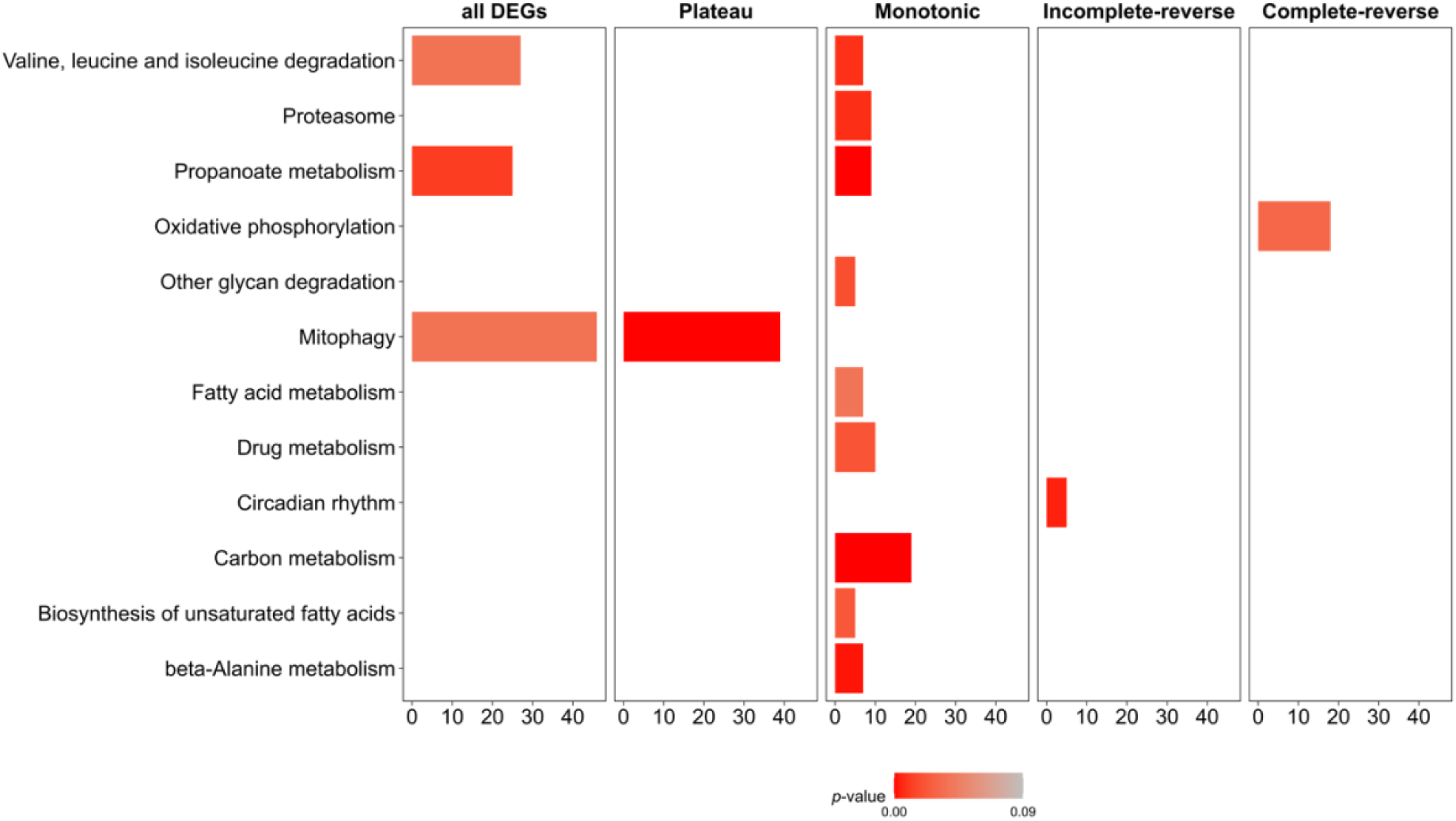
Enrichment of Kyoto Encyclopedia of Genes and Genomes (KEGG) pathways across differentially expressed genes with conserved co-expression patterns. The significant pathways are shown for each group. The ‘all DEGs’ group refers to all the differentially expressed genes identified in generations 7 and/or 31. The pathway descriptions are shown on the y axis, and the -log_10_-transformed *p*-values of the enrichment test are shown on the x axis. For the complete list of pathways and associated *p*-values see Table S2.

**Fig. S7.**
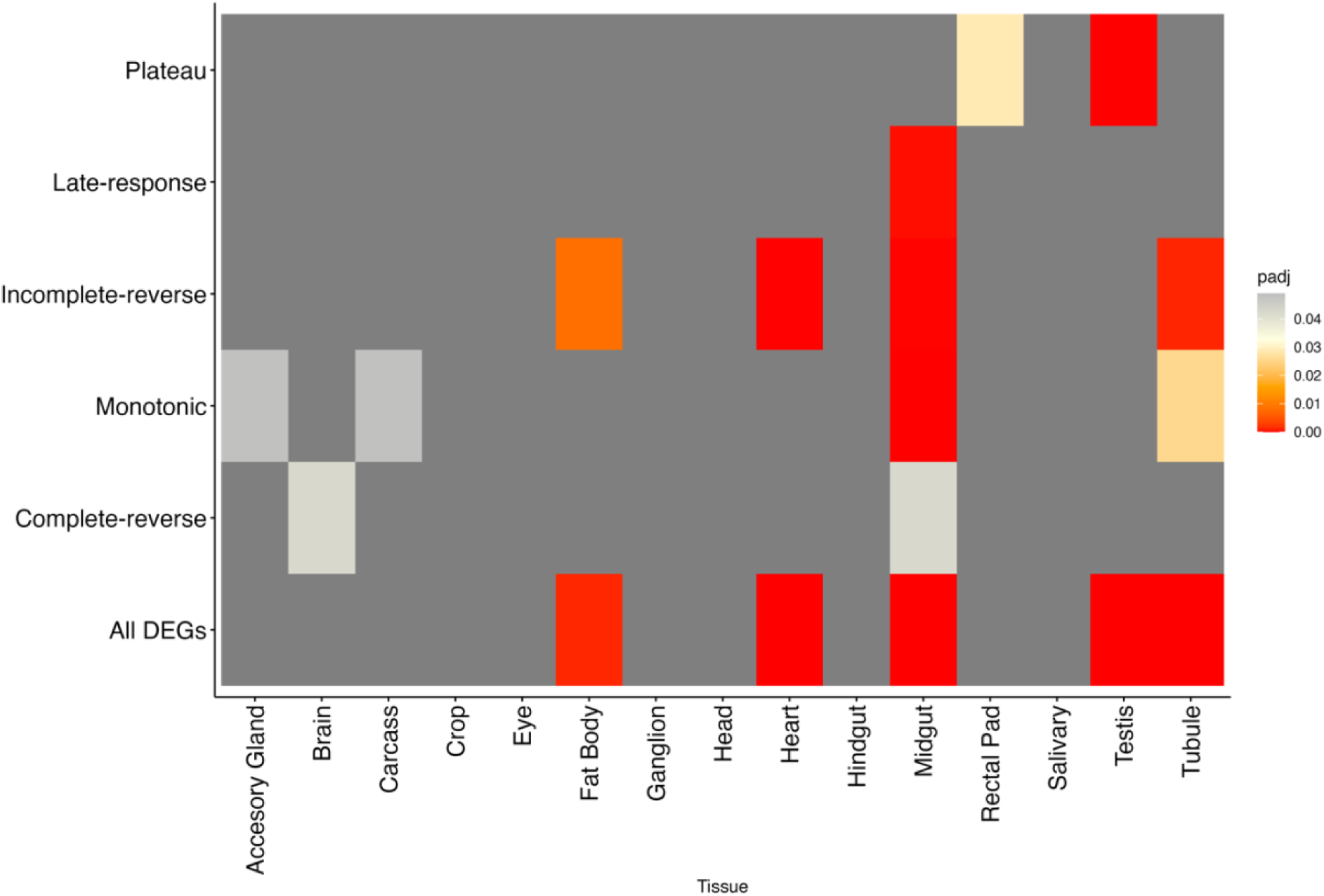
Tissue enrichment across differentially expressed genes with conserved co-expression patterns. The ‘all DEGs’ group refers to all the differentially expressed genes identified in generations 7 and/or 31. The transcriptomic groups are shown on the y axis, and the tested tissues on the x axis. The significance levels are depicted in different colors. Note that dark grey corresponds to the non-significant tissues. For the complete list of tissues and associated *p*-values see Table S2.

**Fig S8.**
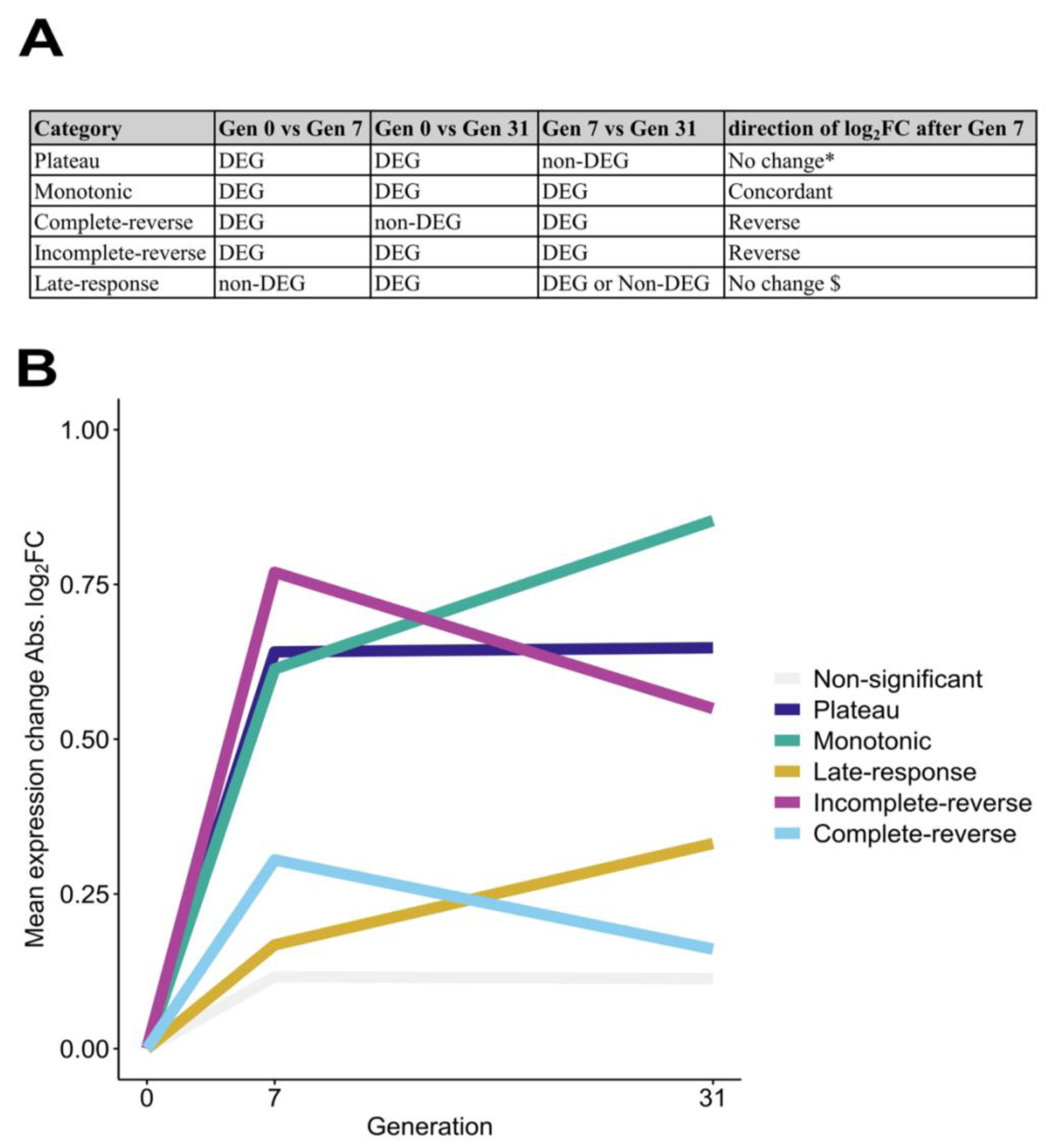
Classification of differentially expressed genes (DEGs) based on the patterns of co-expression. A) The identified DEGs were classified into 5 groups based on the significance and direction of expression change (log_2_FC). DEGs were identified by comparing Gen 0 vs Gen 7, Gen 0 vs Gen 31, and Gen 7 vs Gen 31, and *p*-values across the three contrasts were corrected for multiple testing using false discovery rate (FDR) method. The log_2_FC was used to classify the genes with a change in the direction of expression, e.g. concordant and reverse. * These genes are not differentially expressed between generations 7 and 31. $ These genes are not differentially expressed between generations 0 and 7. 1) The ‘plateau’ group contained genes that their expressions significantly changed in both evolved time points, CGE-7 and CGE-31, compared to the founder population, CGE-0 (p value < 0.05), but no significant change was observed between two evolved time points, CGE-7 and CGE-31 (p value > 0.05). 2) Genes in the ‘monotonic’ group, similar to the ‘plateau’ genes showed significant expression change in both evolved time points, CGE-7 and CGE-31, compared to the founder population, CGE-0. But unlike the ‘plateau’ genes, the expression between CGE-7 and CGE-31 was significant. The direction of change in the expression of these genes (either decreasing or increasing) was consistent across all time intervals. 3) Genes in the ‘complete-reverse’ group had significant expression changes in CGE-7 compared to CGE-0. After CGE-7, the expression level reverted back to the level observed in CGE-0. 4) The gene expression patterns in the ‘incomplete-reverse’ group were similar to the ‘complete-reverse’ group. However, the expression changes between the last time point (CGE-31) and the founder population (CGE-0) were significant (p-value < 0.05). 5) Genes in the ‘late-response’ group had significant expression changes in the last time point (CGE-31) compared to the founder population (CGE-0). However, the expression of these genes was not significant in the first time point (CGE-7) compared to the founder population (CGE-0). B) The mean gene expression change (mean absolute log_2_FC) of the co-expressed gene groups, was calculated at generations 7 and 31 and plotted against generation 0, which was set as common intercept to draw the trajectories. Note that non-significant genes show the lowest log_2_FC.

**Fig. S9.**
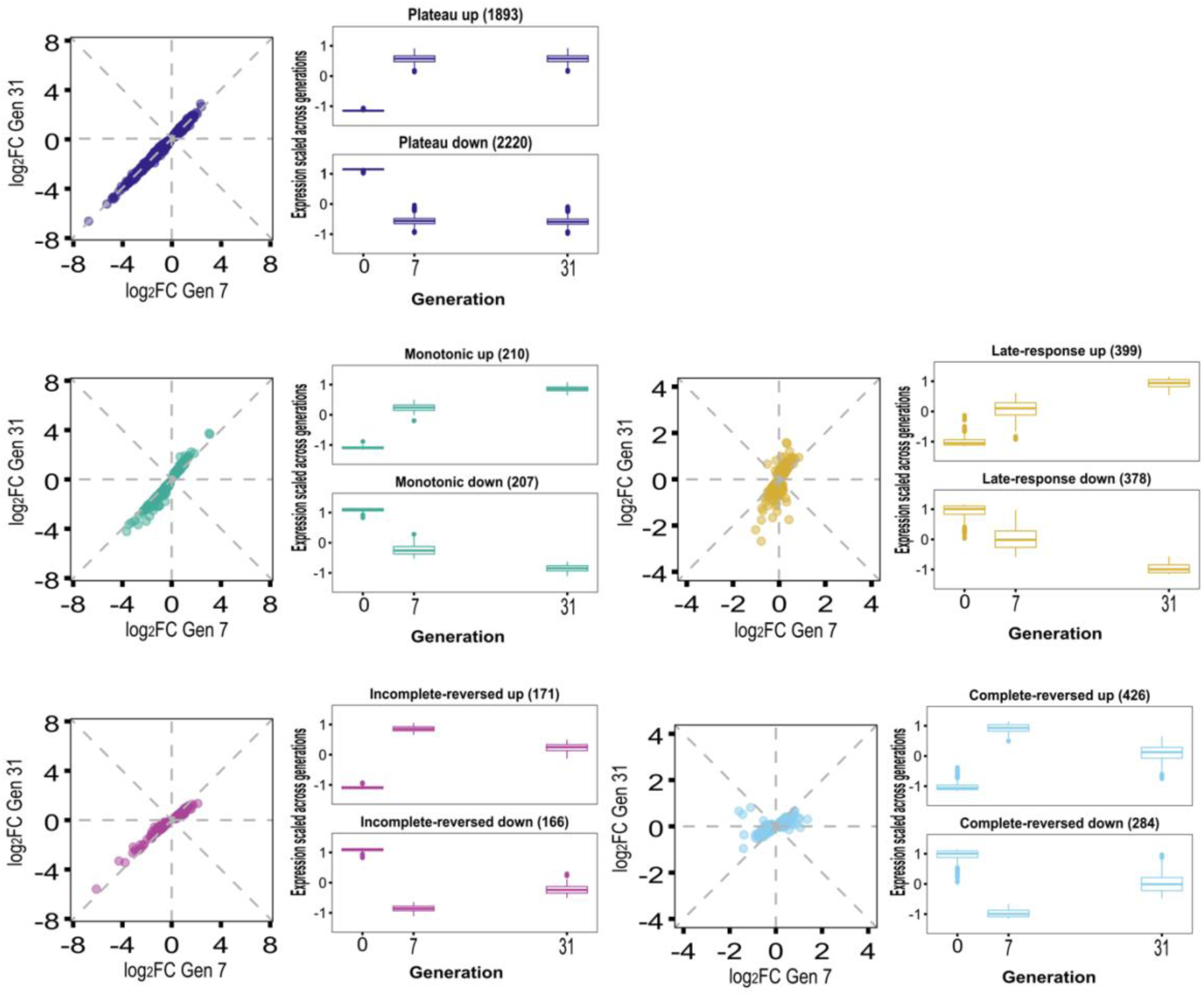
Temporal dynamics of differentially expressed genes (DEGs) with conserved co-expression patterns. For each group, the dot plot shows the evolutionary responses of the evolved populations, quantified as the log_2_ fold-change (log_2_FC) gene expression at generation 7 (x axis) and 31 (y axis) compared to the founder population. The boxplot for each group shows the scaled and centered mean expression (log_2_CPM) of genes across time for up- and down-regulated genes. This plot provides complementary support for the classification of DEGs into groups with conserved co-expression patterns, as shown in Fig. 3 and S8. The number of DEGs in each group is indicated in parentheses.

**Fig. S10.**
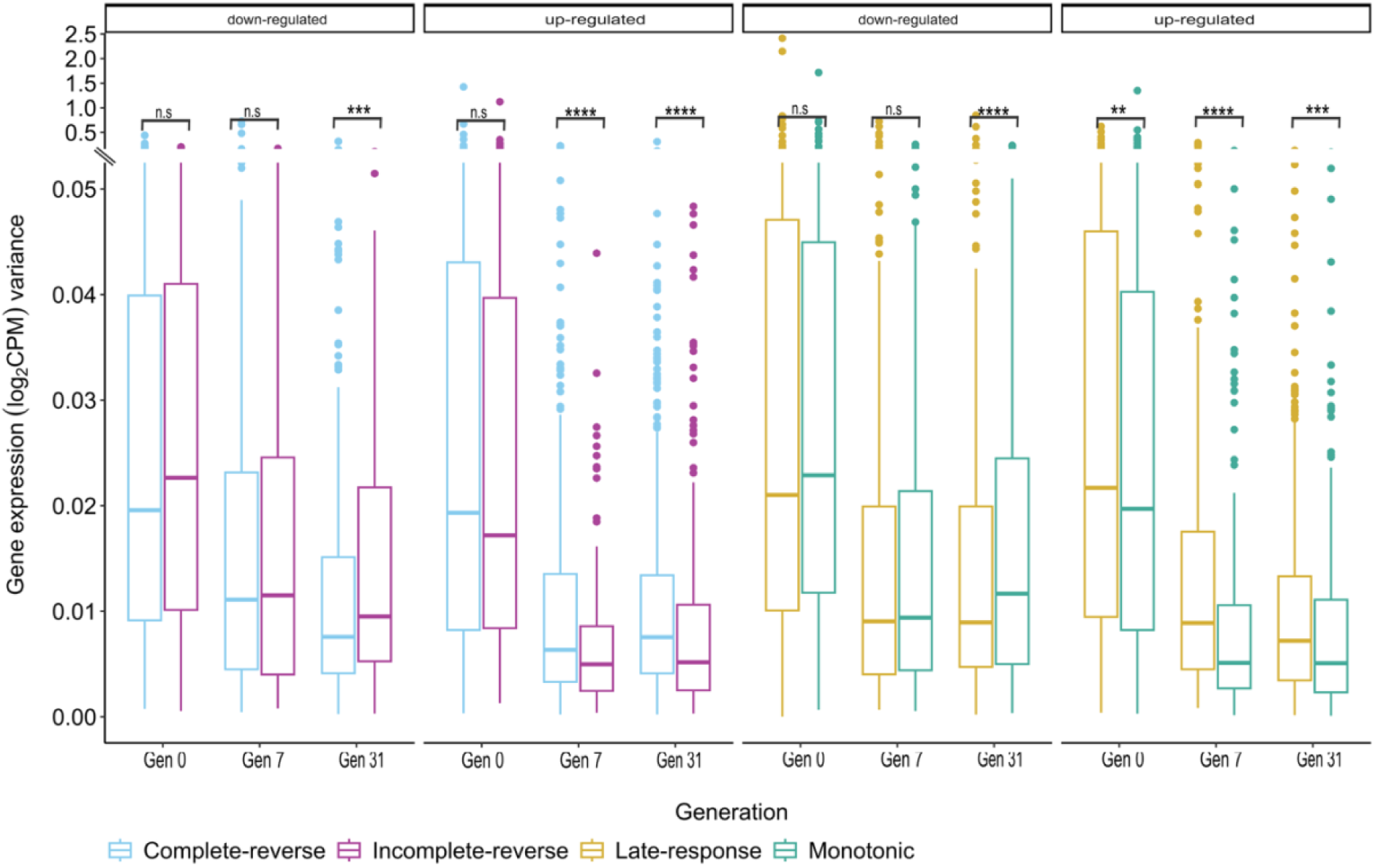
Gene expression variance between transcriptomic groups. The median expression variance of down-, and up-regulated differentially expressed genes differs between the ‘late-response’ and ‘monotonic’ groups, as well as between the ‘complete-reverse’ and ‘incomplete-reverse’ groups (in most cases), suggesting that significant differences distinguish these groups as unique. For each gene the expression variance is computed across 6 evolved replicate populations in each generation. Wilcoxon rank sum tests were performed (see Table S3 for all statistics test results).

**Fig. S11.**
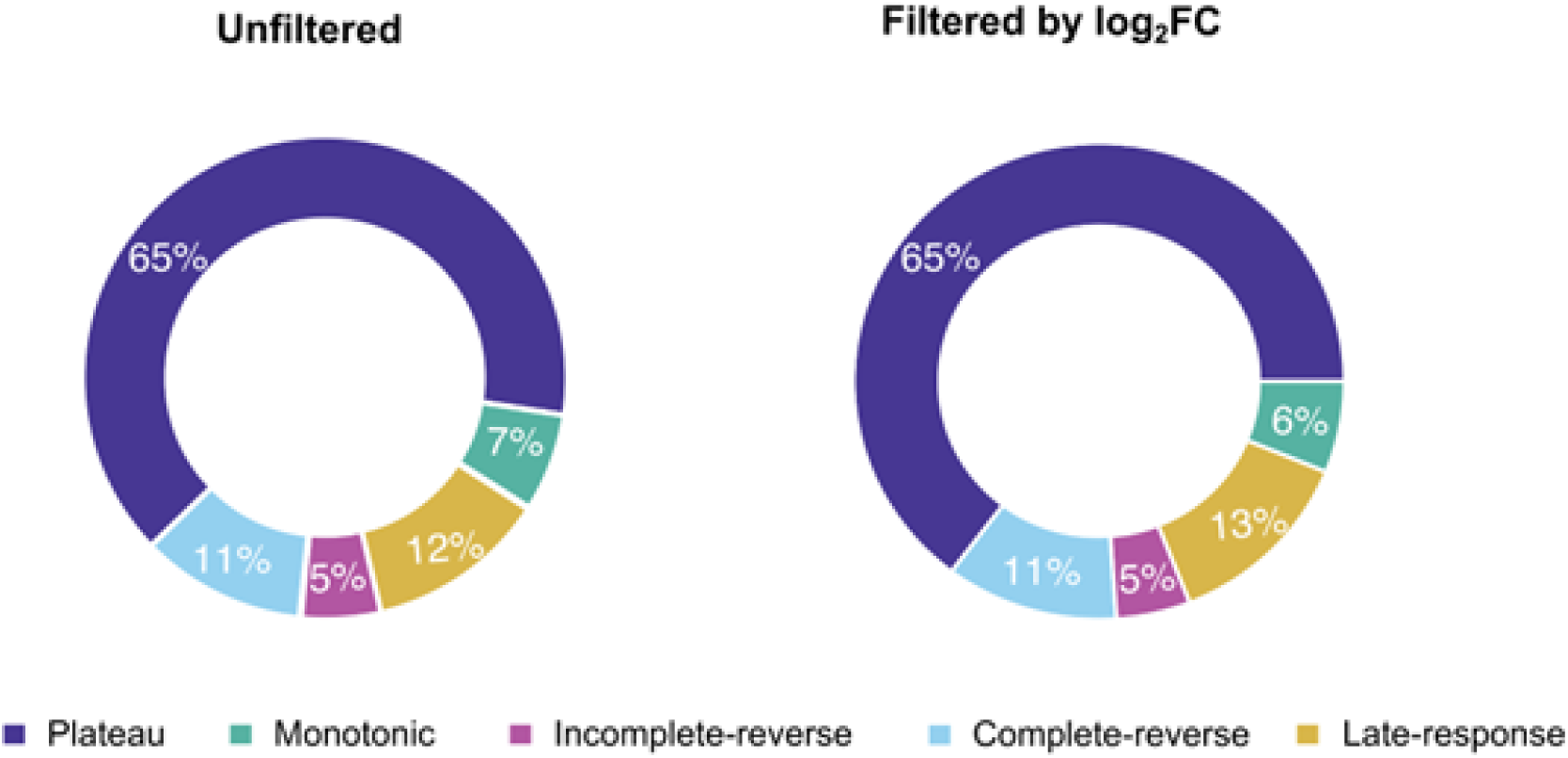
Classification of differentially expressed genes based on the patterns of co-expression is robust to filtering the gene with low magnitude of expression change. Only differentially expressed genes with absolute expression change higher than 0.32 (|log_2_FC| > 0.32) were classified into groups with conserved patterns of co-expression. The proportion of groups remained the same after filtering by log_2_FC, which suggests that the observed pattern is not driven by genes with small changes in expression.

**Fig. S12.**
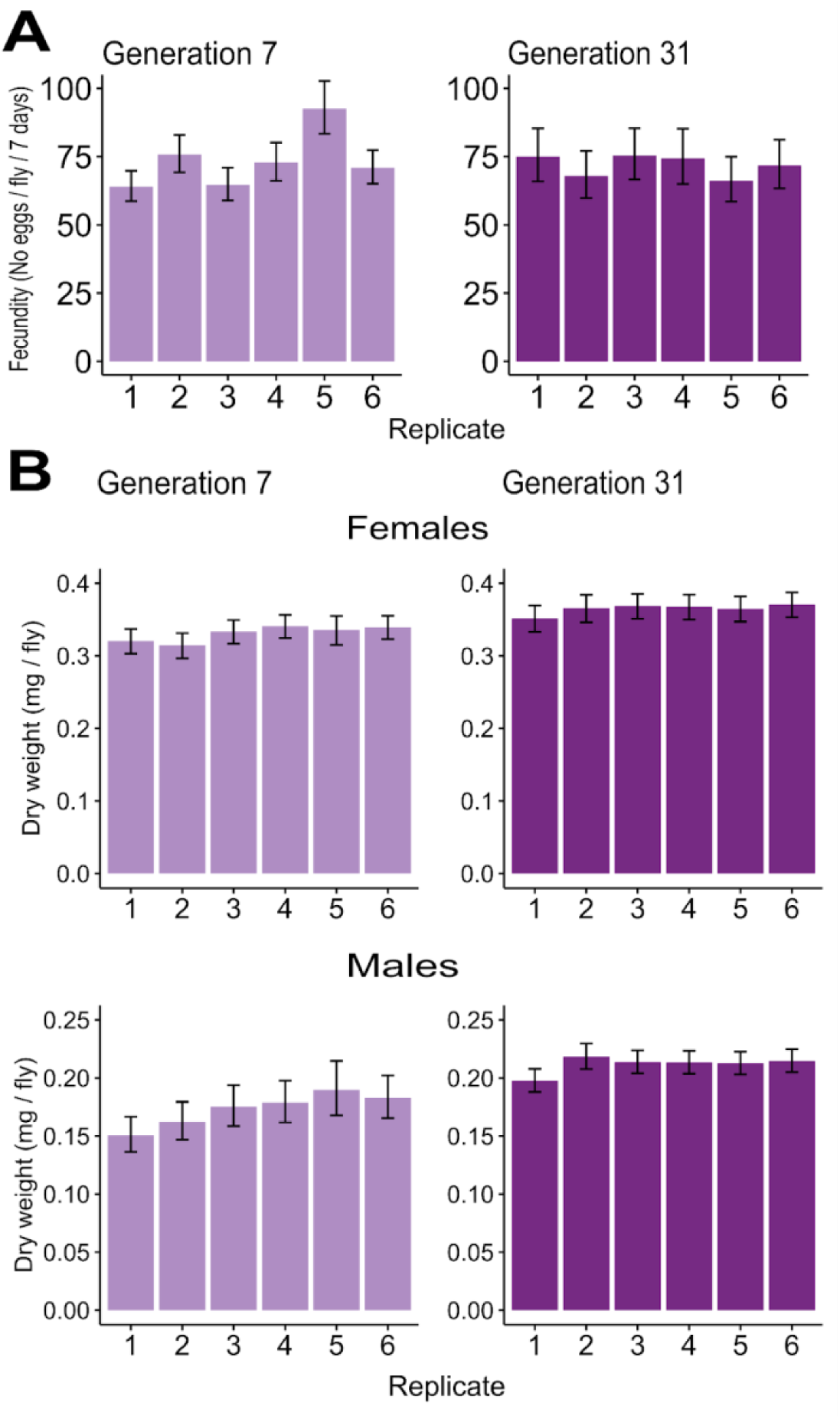
Convergence of fecundity and weight among replicate populations. A) Convergent evolution of fecundity occurred at generation 31 as there was no significant difference among the replicates. However, replicate 5 was significantly more fecund than all other replicates at generation 7, and replicate 2 was also more fecund than replicate 1 at generation 7 (*p*-value < 0.05). Fecundity is presented as the average number of eggs laid by a female during 7 days. B) The replicate populations (females and males) converged for the dry body weight at generations 7 and 31 (p-value > 0.05). Significance was assessed using *emmeans* with option *pairwise* option. *P*-values were corrected across all contrasts within phenotype using the Benjamini-Hochberg’s false discovery (FDR) method (Benjamini & Hochberg, 1995) with an alpha threshold at 0.05. The bars depict the predicted means of the linear model, and error bars show the 95% confidence intervals of the least-square means. See Table S4 for all test results.

**Fig. S13.**
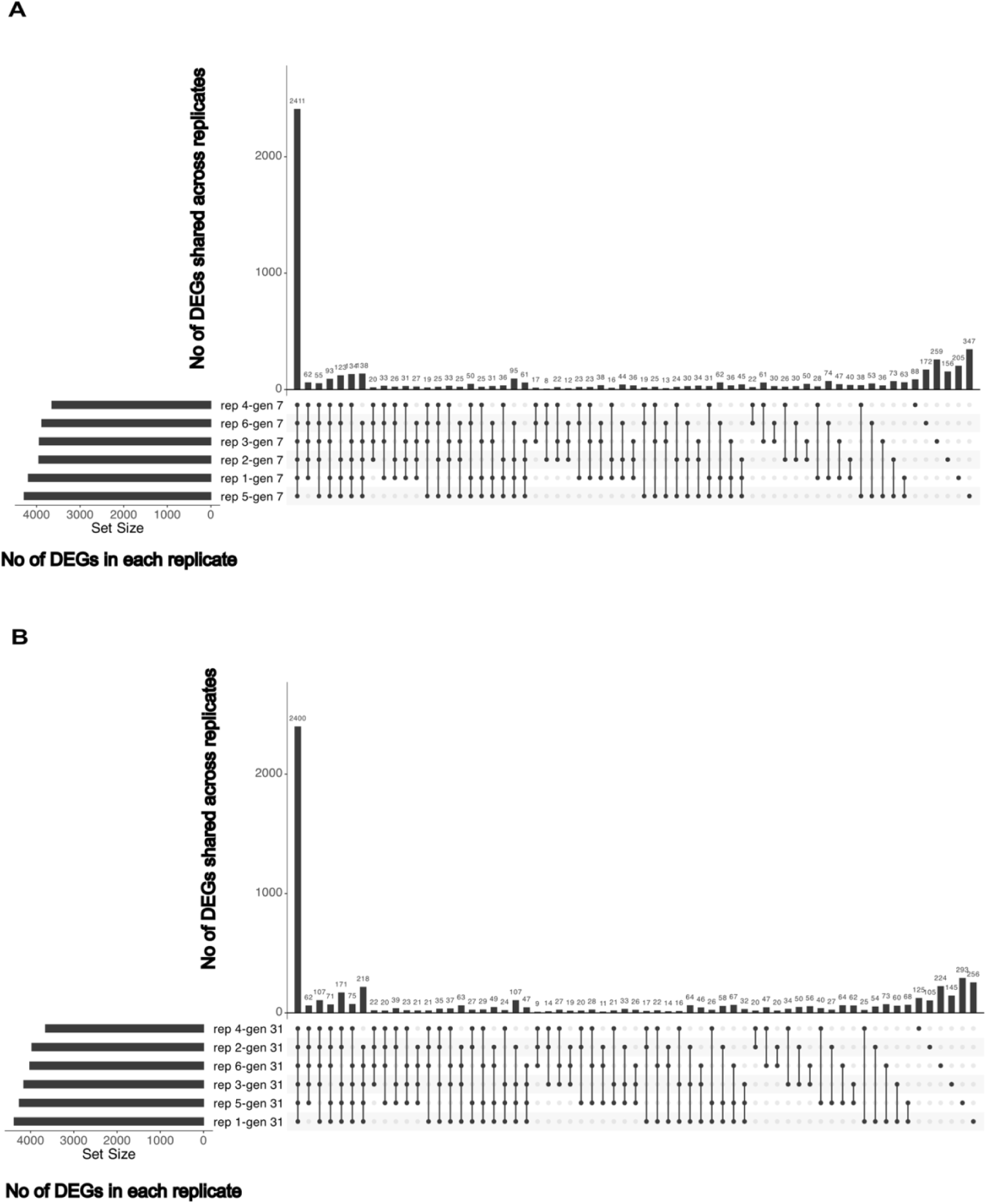
The number of differentially expressed genes (DEGs) shared across different evolved replicate populations at generation 7 (A) and 31 (B). The vertical lines connect the replicate populations, depicted by dots, that have shared DEGs. The majority of DEGs are shared by all six replicates in both generations. rep: replicate, gen 7: generation 7, gen 31: generation 31.

**Fig. S14.**
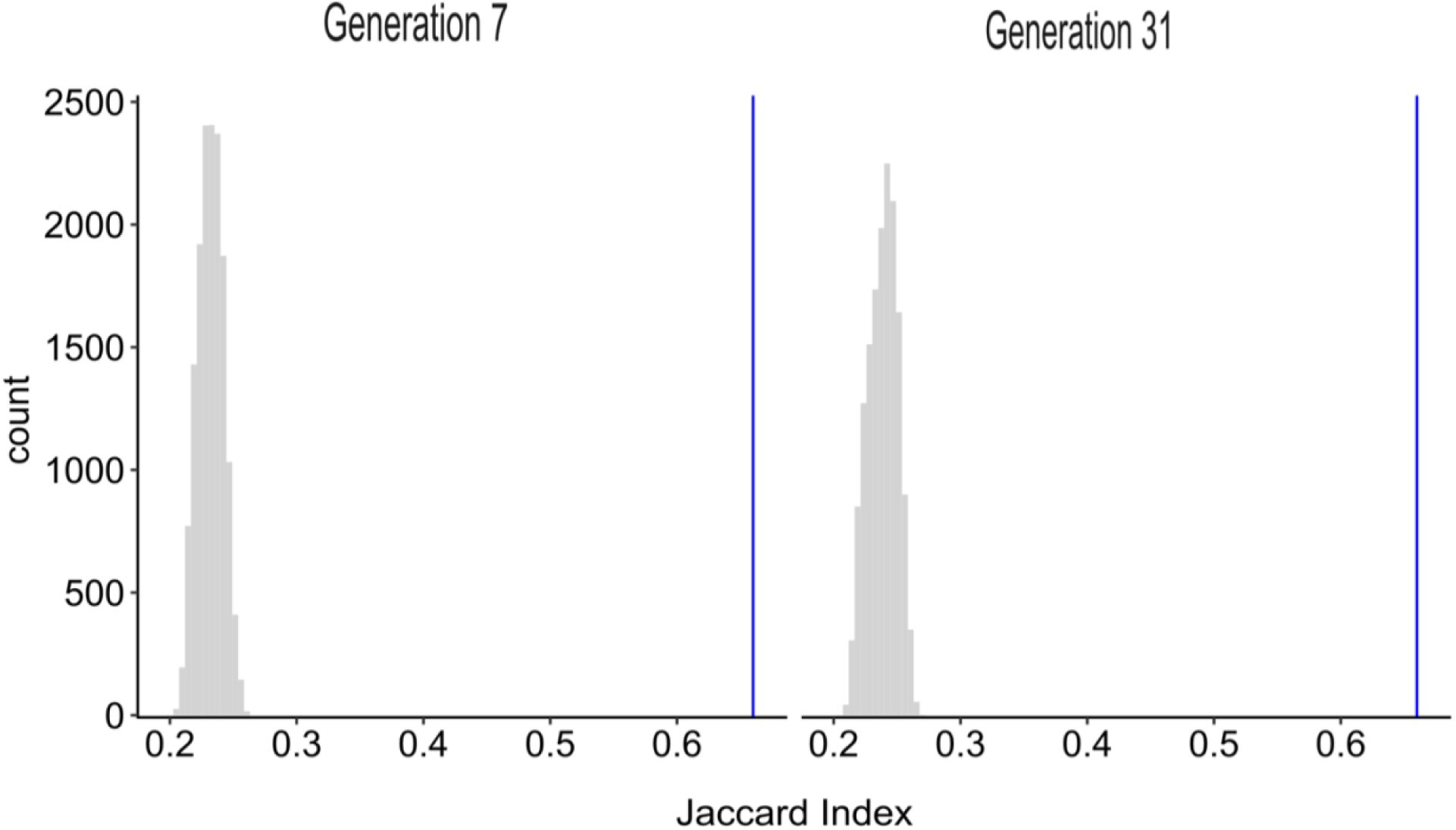
The average Jaccard index among replicate populations is significantly higher than the null distribution. The null distribution (grey) for the Jaccard index was generated using delete-d jackknifing with 1000 iterations. In each iteration, DEGs in each replicate were randomly selected from the total of expressed genes in the transcriptome. Samples sizes (number of genes) matched the observed DEGs for each replicate in each time point. The null distribution showed that random sampling produced significantly lower parallelism than the empirical Jaccard index (vertical blue line).

**Fig. S15.**
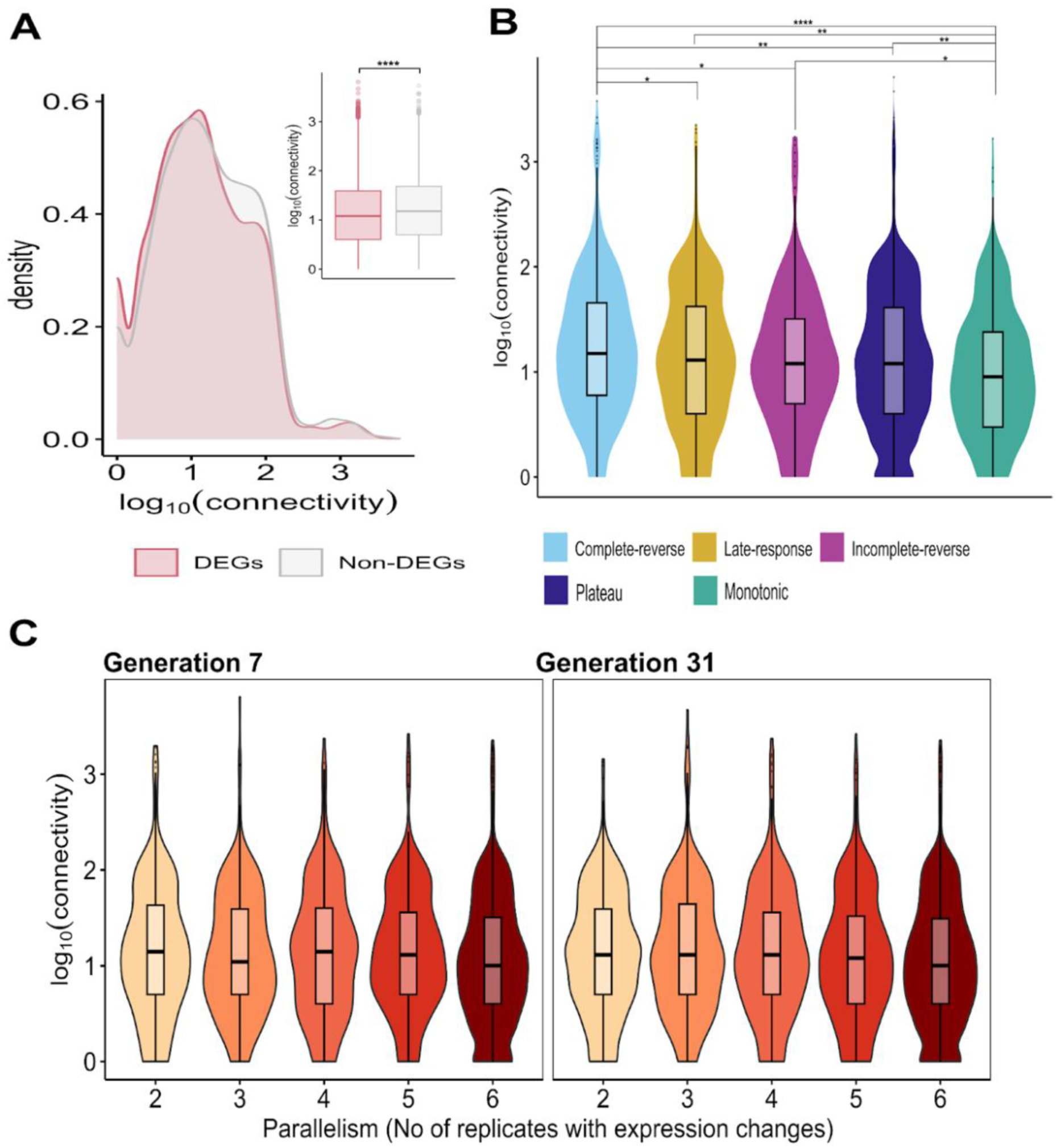
Low-to-intermediate pleiotropic genes facilitate adaptation and drive parallel changes among replicate populations. A) Pleiotropy levels (log_10_-transformed gene connectivity in a gene regulatory network) of differentially expressed genes (DEGs) and non-differentially expressed genes (Non-DEGs) detected at generations 7 and/or 31 with available connectivity data (5,577 DEGs, and 3,369 non-DEGs). B) Pleiotropy levels of the DEGs groups with conserved patterns of co-expression. Two-tailed Wilcoxon rank-sum tests were conducted for two groups in (A) and all pairwise comparisons in (B). Benjamini-Hochberg’s false discovery (FDR) method (Benjamini & Hochberg, 1995) was applied to correct the *p*-values across all the test in A and B, and significant results are indicated: (*) *p*-value < 0.05, (**) *p*-value < 0.01, (***) *p*-value <0.001, and (****) *p*-value <0.0001 (see Table S5 for all statistical test results). C) Pleiotropy levels (log_10_ gene connectivity) of genes with differential expression change across different numbers of replicate populations (e.g., group 2: DEGs shared by 2 replicates). Note that group 6 contains genes with the lowest level of parallelism. Two-tailed Wilcoxon rank-sum tests were conducted for all pairwise comparisons in each generation. FDR correction was applied across all tests. Genes with the highest level of parallelism (group 6) are significantly less pleiotropic than groups 2, 4 and 5 of parallelism at generation 7, and than groups 2, 3, and 4 at generation 31 (see Table S7 for all statistical test results).

**Fig. S16.**
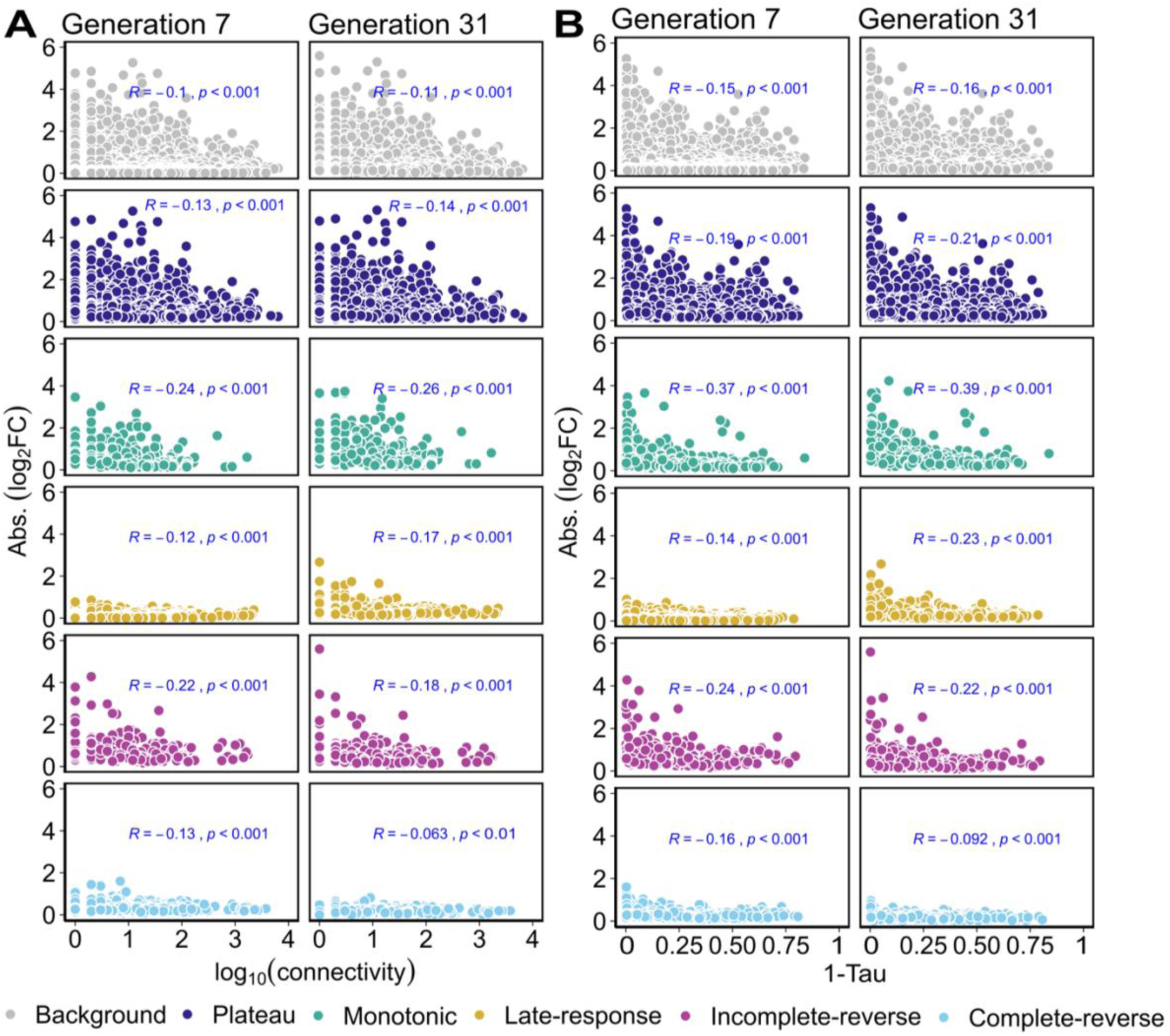
The magnitude of gene expression is negatively correlated with the levels of pleiotropy. The transcriptomic response expressed as the absolute log_2_ fold-change (log_2_FC) between the founder and evolved populations in each generation is shown on the y axis, and the levels of pleiotropy expressed as log_10_ gene connectivity (A), and tissue specificity (1-Tau) (B) are shown on the x axis. Kendall’s tau rank correlation between the absolute expression changes and pleiotropy was computed for each transcriptomic group and generation. ‘Background’ refers to all differentially expressed genes. Note that the strongest correlation is for the ‘monotonic’ group (R = -0.37 at generation 7, R = -0.39 at generation 31). Kendall’s correlation coefficients (R) and significance (*p*) are depicted in blue within each transcriptomic group. Benjamini-Hochberg’s false discovery (FDR) method (Benjamini & Hochberg, 1995) was applied to correct the *p*-values across all tests within a pleiotropy metric.

**Fig. S17.**
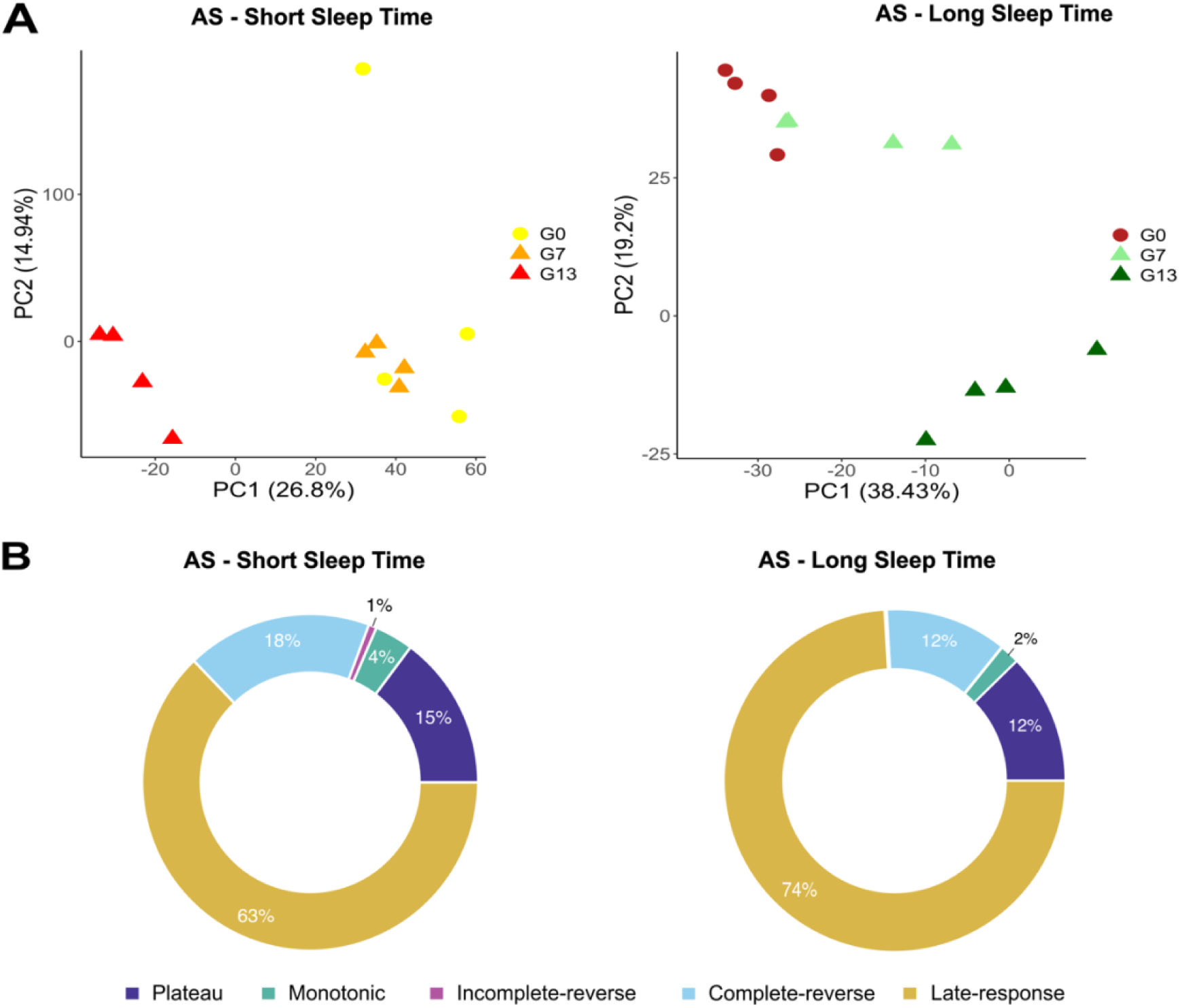
Differentially expressed genes (DEGs) with conserved co-expression patterns in populations under artificial selection. Replicate populations of *Drosophila melanogaster* were selected for longer and shorter sleep duration for 13 generations (Souto-Maior et al., 2023). The focal phenotypes, i.e. sleep duration, did not plateau during the experiment. A) Principal component analysis (PCA) using all expressed genes after filtering the lowly expressed genes (12637 and 12457 genes in the ‘Short Sleep Time’ and ‘Long Sleep Time’, respectively) showed divergence between the evolved populations in different generations. B) A total of 3,108 and 2,658 DEGs were identified in the ‘Short Sleep Time’ and ‘Long Sleep Time’ experiments, respectively. The DEGs between generations 0 and 7, and between generations 7 and 13 were classified into groups with conserved co-expression patterns similar to our study. The pie chart illustrates the proportion of DEGs in each group. Only a small proportion of DEGs were in the ‘plateau’ group (12% and 15% of DEGs in the ‘Short Sleep Time’ and ‘Long Sleep Time’, respectively).

**Fig. S18.**
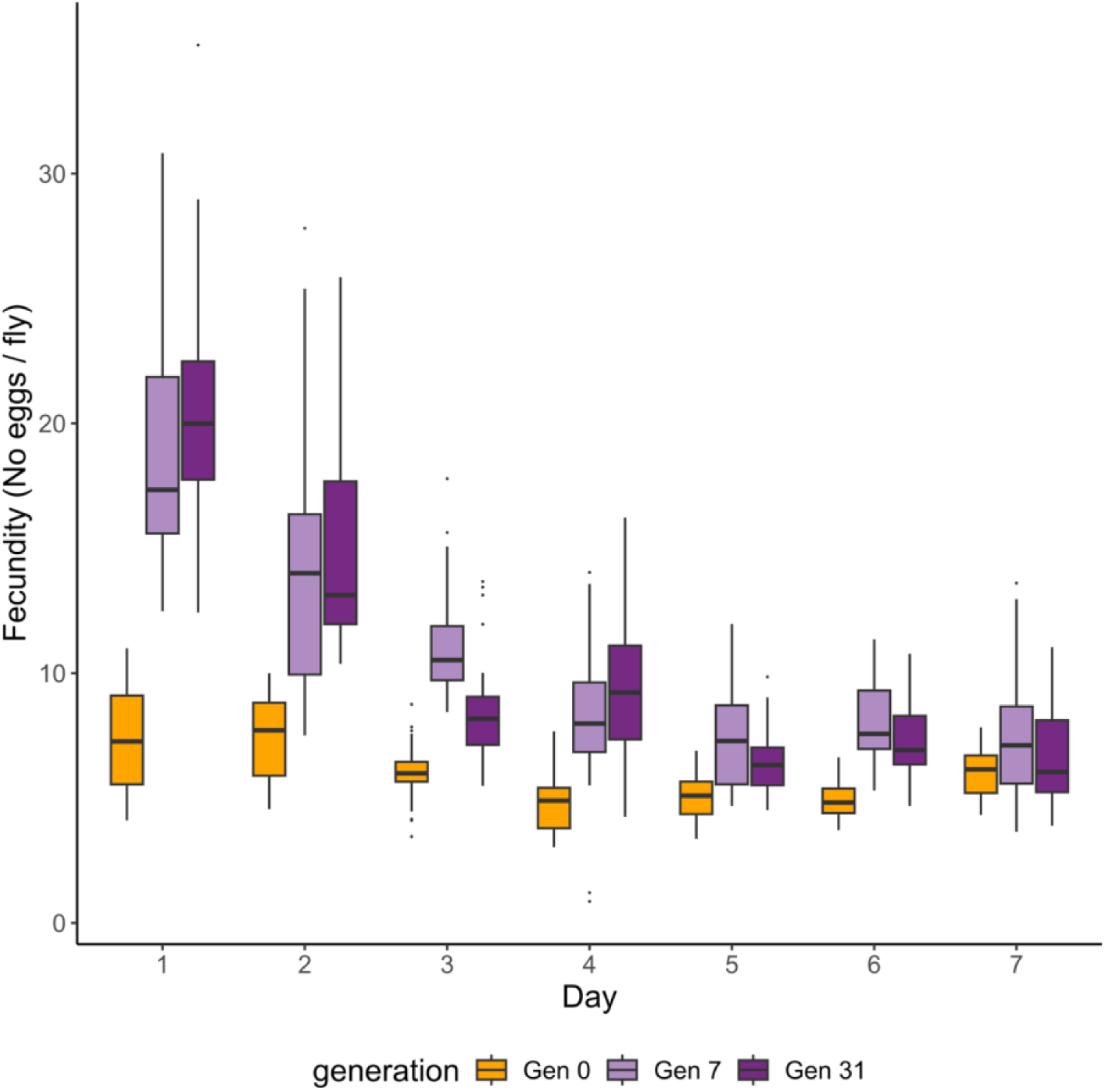
Daily rate of fecundity in the founder and evolved populations. The daily number of eggs/female in sub-replicates are shown. In Gen 0, we measured fecundity in 10 replicates of the founder population (each with 3 sub-replicates), thus a total of 30 sub-replicates. In Gen 7 and Gen 31, fecundity was measured in 3 sub-replicates for each of the 6 evolved populations, a total of 18 sub-replicates. The evolved populations had the highest fecundity on day 1: mean = 18 eggs/day at Gen 7, mean = 20 eggs/day at Gen 31. The maximum number of eggs/day also occurred on day one: mean = 30 eggs/day at Gen 7, mean = 35 eggs/day at Gen 31.

**Fig. S19.**
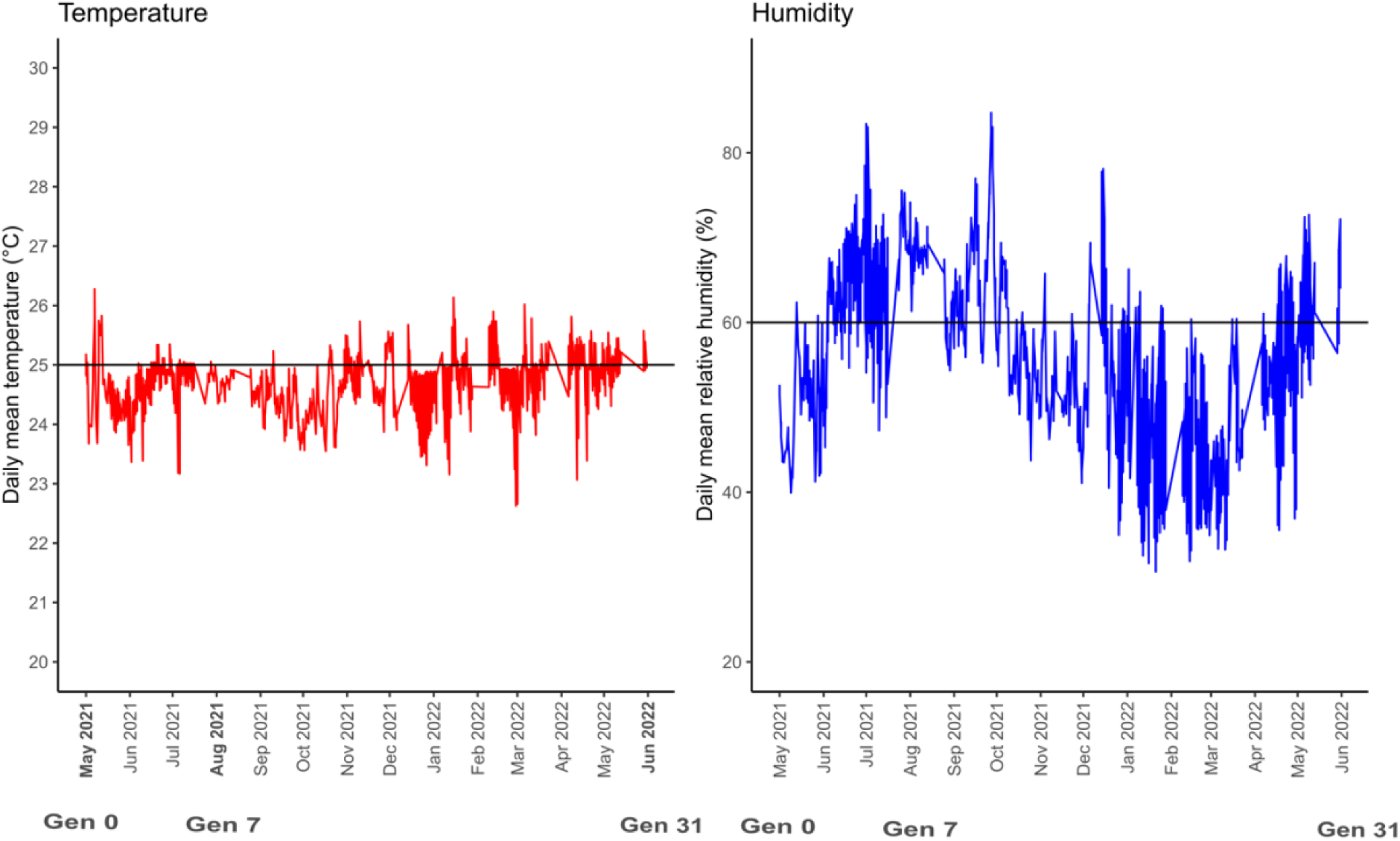
The daily records of temperature and humidity during the evolution experiment. Four sensors in the walk-in incubator room collected temperature and humidity data every minute. The daily average temperature and humidity data across four sensors are shown. Time in months is shown on the x axis, and the generations (Gen 0, Gen 7 and Gen 31) where samples were collected from the experimental cages for the common garden experiments are marked in bold. During the common garden experiments the populations were maintained in incubators to ensure even more precise control of temperature and humidity.

**Fig. S20.**
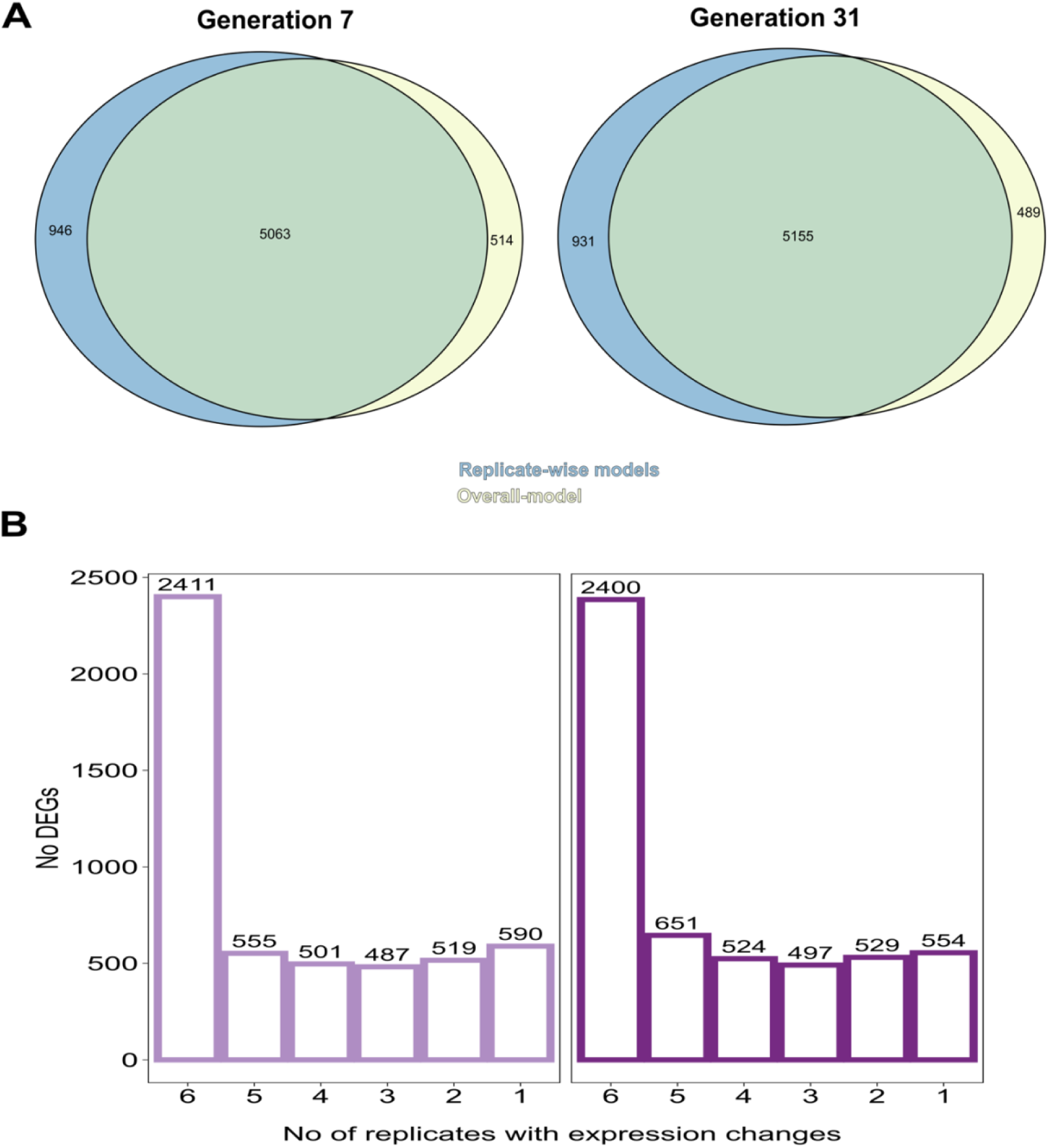
Overlap of the differentially expressed genes (DEGs) identified using the overall and replicate-specific models. (A) Most of the DEGs detected by the overall model (which compares all six replicate populations together to the founder population for the identification of DEGs) were also detected by the replicate-specific models (which identified the gene expression change by comparing each replicate population separately to the founder population) in generation 7 or 31. The overall model is more conservative than the replicate-specific model as it identifies fewer DEGs (432 and 442 fewer DEGs in generations 7 and 31, respectively). (B) Distribution of the levels of parallelism across genes detected by both the overall and replicate-specific models (intersection of the overall and replicate-specific gene sets in A). Most of the identified DEGs by the overall model and the replicate-specific models were shared (2411 and 2400 genes in generations 7 and 31, respectively). A small fraction of the DEGs (<12%, 590 and 554 genes in generations 7 and 31, respectively) were specific to one replicate only either in generation 7 or 31.

